# TNF and type I IFN induction of the IRG1-itaconate pathway restricts *Coxiella burnetii* replication within mouse macrophages

**DOI:** 10.1101/2023.07.07.548079

**Authors:** Mark A. Boyer, Natasha Lopes Fischer, Sunny Shin

**Affiliations:** Department of Microbiology, University of Pennsylvania Perelman School of Medicine, Philadelphia, PA 19104

## Abstract

The intracellular Gram-negative bacterium *Coxiella burnetii* replicates within macrophages and causes a zoonotic disease known as Q fever. In murine macrophages, the cytokine tumor necrosis factor (TNF) is critical for restriction of intracellular *C. burnetii* replication. Here, we show that TNF collaborates with type I interferon (IFN) signaling for maximal control of *C. burnetii*. We found that TNF and type I IFN upregulate the expression of the metabolic enzyme immune responsive gene 1 (IRG1), also known as cis-aconitate decarboxylase 1 (ACOD1), and that IRG1 is required to restrict *C. burnetii* T4SS translocation and replication within macrophages. Further, we show that itaconic acid, the metabolic product of IRG1, restricts *C. burnetii* replication both intracellularly and in axenic culture. These data reveal that TNF and type I IFN upregulate the IRG1-itaconate pathway to restrict intracellular *C*. *burnetii* replication within murine macrophages.

## Introduction

The Gram-negative facultative intracellular bacterium *Coxiella burnetii* is the causative agent of the zoonotic disease known as Q fever (1). Often asymptomatic, Q fever presents as an acute atypical pneumonia, and can develop into a long-term chronic infection that can lead to hepatitis or endocarditis, with an increased risk of severe disease to individuals with immunodeficiencies or heart issues and pregnant women (2, 3). Q fever occurs worldwide and is a significant public health concern with periodic outbreaks, such as one that occurred in the Netherlands from 2007-2010 with over 4,000 cases (4-9). *C. burnetii* is classified as a category B pathogen due to its aerosol mode of transmission and environmental stability (10). Despite this potentially severe threat to human health, the mechanisms underlying how *C. burnetii* interacts with host cells and the critical host factors that restrict bacterial replication remain poorly understood.

The predominant reservoir for *C. burnetii* is domesticated livestock, and the bacteria can easily be transmitted to human hosts via aerosolized secretions (9). Upon inhalation into the lung, *C. burnetii* is taken up primarily by resident alveolar macrophages (11, 12). Following internalization, *C. burnetii* engages both the endolysomal and autophagic pathways (13-15) to form its replicative compartment known as the *Coxiella*-containing vacuole or CCV (16). Full acidification and maturation of this phagolysosome is necessary to stimulate bacterial metabolism and to activate the *C. burnetii* Type IVb Secretion System (T4SS), which delivers over 100 effector proteins to manipulate the host cell (17-19). The T4SS is required for replication as well as maintenance and expansion of the CCV (20-22).

Macrophages employ pattern recognition receptors (PRRs), such as Toll-like receptors (TLRs), to detect pathogen-associated molecular patterns (PAMPs) (23, 24). In particular, TLR2 signaling is important for controlling the intracellular replication of *C. burnetii* within murine macrophages (25, 26). We previously found that TLR2-driven restriction of *C. burnetii* within macrophages was dependent on production of the pro-inflammatory cytokine Tumor Necrosis Factor (TNF) (27). In other contexts, TNF can promote a variety of immune responses, including transcription of inflammatory mediators and induction of cell death (28-30). TNF binding to its receptors TNFR1 and TNFR2 can trigger a signaling cascade that upregulates many genes and antimicrobial programs (31), but can also be a key driver of pathologic inflammation. Thus, anti-TNF therapy is a frequent treatment for autoimmune diseases, such as rheumatoid arthritis, Crohn’s disease, and multiple sclerosis. However, patients on TNF blockade are more susceptible to intracellular bacterial pathogens, including *Mycobacterium, Listeria*, *Legionella*, *Salmonella*, and *Coxiella* (32-34). Surprisingly, the mechanisms underlying how TNF restricts *C. burnetii* and other intracellular bacterial pathogens are not fully understood.

Here, we find that maximal TNF-dependent restriction of *C. burnetii* replication relies on basal type I interferon (IFN) signaling in murine macrophages. Furthermore, we show that TNF and IFN collaborate to induce maximal expression of the host gene Immune Responsive Gene 1 (*Irg1*). *Irg1* encodes a cis-aconitate decarboxylase (ACOD1) that converts the TCA intermediate aconitate into itaconate, which possesses antimicrobial activity against intracellular pathogens including *L. pneumophila, M. tuberculosis, and Salmonella* (35-37) Our data indicate that IRG1 mediates TNF and IFN-mediated restriction of *C. burnetii* T4SS translocation and replication within murine macrophages. Addition of itaconate to *Irg1^-/-^* cells restored inhibition of *C. burnetii* T4SS secretion and intracellular replication. Furthermore, itaconate was sufficient to inhibit bacterial replication in axenic cultures. Overall, our findings reveal that TNF and type I IFN signaling collaborate to induce IRG1 expression, and that IRG1 is required for TNF and type I IFN-dependent restriction of *C. burnetii* in murine macrophages. Collectively, these findings provide important insight into the mechanisms underlying how TNF restricts *C. burnetii* replication within murine macrophages.

## Materials and Methods

### Bacterial culture conditions

All experiments were conducted using variants of the avirulent Nine Mile Phase II (NMII) strain of *Coxiella burnetii* (RSA 439, Clone 4): wild-type (WT) *C. burnetii*, WT *C. burnetii* expressing mCherry (38), an *icmL:Tn* strain with a non-functional T4SS due to a transposon insertion in the *icmL* gene (20), and WT and *icmL:Tn* strains harboring a plasmid encoding *blaM* alone or *blaM* fused to the T4SS effector gene *CBU_0077* (20). The bacteria were cultured in acidified citrate cysteine medium 2 (ACCM-2; Sunshine Scientific Products) (39) at a starting concentration of 2×10^6^ bacteria/ml for 7 days at 37°C and 5% CO_2_ and 2.5% O_2_. One day after inoculation, kanamycin (275 mg/ml) was added to the icmL:Tn cultures and chloramphenicol (3 mg/ml) was added to the cultures of the strains containing the BlaM- or BlaM-0077-expressing plasmids. After 7 days, the cultures were quantified by isolating the genomic DNA from 1ml of inoculate using GenElute Bacterial Genomic DNA kit (Sigma) and measuring the genomic equivalents (GE) by quantitative PCR (qPCR) using primers for *C. burnetii dotA* gene (FWD: GCGCAATACGCTCAATCACA and REV: CCATGGCCCCAATTCTCT). The plasmid pGEM-T encoding the *C. burnetii dotA* gene was serially diluted and used to generate a standard curve for qPCR to enable the calculation *C*. *burnetii* genome equivalents. For growth assays of *C. burnetii* axenic cultures, pharmacological agents were added to the ACCM at the time of inoculation or at a time otherwise specified.

### Mouse strains

For generation of murine bone marrow-derived macrophages, the following mouse strains were used. C57BL/6J (strain 000664), C57BL/6NJ (strain 005304), *Tnf^-/-^*(strain 005540), *Tnfr1^-/-^* (003242), *Tnfr2^-/-^* (strain 002620), *Nos2^-/-^* (strain 002609), *Gp91phox^-/-^* (strain 002365) and *Irg1^-/-^* mice (strain 029340) were purchased from Jackson Laboratory. *Ripk3^-/-^, Casp8^-/-^Ripk3^-/-^,* and *Ripk1^kd/kd^* mice were generously provided by Dr. Igor Brodsky, and *Ifnar^-/-^*mice were generously provided by Dr. Carolina Lopez. All animals were housed and bred in specific pathogen-free conditions in accordance with the Animal Welfare Act (AWA) and the guidelines of the University of Pennsylvania Institutional Animal Use and Care Committee. All protocols in this study were approved by the University of Pennsylvania Institutional Animal Use and Care Committee (protocol #804928).

### Cell culture

Murine bone marrow-derived macrophages (BMDM) were generated by harvesting bone marrow cells from the bones of 8-12 week old mice. Cells were cultured in 10cm petri dishes with Dulbecco’s modified Eagle’s Medium (DMEM) + GlutaMAX (Invitrogen) supplemented with 20% fetal bovine serum (FBS), 1mM sodium pyruvate (Invitrogen), 20 ng/ml recombinant murine M-CSF (Peprotech), and penicillin/streptomycin (Corning) for 7 days at 37°C and 5% CO_2_. 24 hours prior to infection, BMDMs were harvested using PBS with 2mM EDTA, centrifuged at 1200rpm for 10 minutes, and then resuspended in growth media with only 10% FBS and without antibiotics. BMDMs were seeded into 24-well non-tissue culture-treated plates for flow cytometry experiments or either 24- or 48-well tissue culture-treated plates for all other experiments. All recombinant proteins and pharmacological agents used for *in vitro* stimulation or inhibition were added during replating, one day prior to infection, unless otherwise specified.

Recombinant proteins and pharmacological agents were added to the replating media at specific concentrations, as indicated in the figure legends: murine TNF-α (Peprotech), universal IFN-α (R&D Systems) and itaconic acid (Fisher Scientific). For itaconic acid, media was normalized back to original pH following addition of itaconic acid.

### Infection

For *in vitro* infection, cells were mock-infected with ACCM or infected with *C. burnetii*. Infected culture plates were spun down for 10 minutes at 1200 rpm and incubated for 4 hours to allow for bacterial entry into cells. After 4 hours, media was aspirated from the wells and cells were washed once with PBS. For Day 0 timepoints, samples were harvested immediately after washing with 500μl sterile H_2_O; for all other timepoints, fresh culture media with or without recombinant proteins or pharmacological agents was added back (500μl for 24-well plates, 300μl for 48-well plates) and cells returned to incubation at 37°C and 5% CO_2_. For growth curves, cells were fed once on Day 4 an additional 500μl of culture media. Genomic DNA was isolated from cell lysates using the GenElute Bacterial Genomic DNA kit (Sigma). *C. burnetii* GEs were measured by qPCR using primers for the *dotA* gene, as described in **Bacteria Culture Conditions.**

### siRNA-mediated gene silencing

Silencer Select siRNA oligos were purchased from Life Technologies. Irg1 siRNA s68386 and negative control siRNAs (Negative Control No. 1 siRNA 4390843 and Negative Control No. 2 siRNA 4390846) were utilized at 12 pmol per well. Transfection into BMDMs was performed using Lipofectamine RNAiMax transfection reagent (Invitrogen), following a modified protocol of the “Forward Transfection” protocol from the manufacturer’s manual. Transfection with appropriate siRNAs was performed one day after BMDM replating into 24-well tissue culture-treated plates. 8 hours after transfection, fresh media supplemented with rTNF was added to each well. After 16 hours, cells were then infected as described in the **Infection** section. 72 hours after the initial siRNA transfection, cells were transfected with siRNAs a second time to increase the duration of knockdown. Knockdown efficiency was confirm using qRT-PCR analysis, described below.

### qRT-PCR analysis

To measure RNA expression, RNA was isolated from cells and purified using the RNeasy Plus Kit (Qiagen). First strand cDNA synthesis from purified RNA was performed using SuperScript II reverse transcriptase and Oligo(dT)_12-18_ Primers (Invitrogen). qPCR was performed with the CFX96 real-time system (Bio-Rad) using the SsoFast EvaGreen Supermix with the LOW ROX kit (Bio-Rad). The following primers were used:

*Gapdh:* FWD – AGGTCGGTGTGAACGGATTTG

REV – TGTAGACCATGTAGTTGAGGTCA

*Irg1*: FWD – GGCACAGAAGTGTTCCATAAAGT

REV - GAGGCAGGGCTTCCGATAG

mRNA relative fold induction was calculated using the ΔΔC_T_ method (40), normalizing the cycle threshold (C_T_) of a given gene to that of the housekeeping gene *Gapdh*. To measure siRNA knockdown efficiency, relative mRNA levels of siRNA-treated cells were normalized to control siRNA-treated cells.

### Flow cytometry

For experiments assaying T4SS injection into BMDMs, the *C. burnetii* strains harboring *blaM* or *blaM-0077* plasmids were used, Infected cells were harvested using cold PBS with 2mM EDTA and pelleted at 2500 rpm at 4°C, then resuspended in PBS with 2mM EDTA and 2% BSA. Cells were then loaded with CCF4-AM for 2 hours at room temperature using the LiveBLAzer Fret-B/G Loading with CCF4-AM kit from Thermo Fisher. After 2 hours, cells were fixed with BD Cytofix (BD Biosciences). Fixed cells were then analyzed on a LSR II flow cytometer (BD Biosciences), and the data were further analyzed using FlowJo analysis software (TreeStar).

### Fluorescence microscopy

For microscopic imaging of WT *C. burnetii* expressing mCherry during infection of macrophages, approximately 2.5 x 10^5^ BMDM cells were seeded onto glass coverslips in 24-well plates and infected as described above. To collect samples for imaging, coverslips were washed gently with PBS, fixed in 4% paraformaldehyde in PBS, permeabilized with 0.2% Triton-X, then washed with PBS and mounted on glass slides using Fluoroshield with Dapi (Sigma). Slides were imaged on an Olympus IX81 inverted fluorescence microscope and analyzed and presented using SlideBook software (Intelligent Imaging Innovations, Inc.).

### Statistical analysis

Graphical plotting and statistical analysis were performed using GraphPad Prism software. Statistical significance was determined using either paired, two-tailed Student’s t test or two-way analysis of variance (ANOVA) with Tukey’s test. Differences were considered statistically significant if the P value was < 0.05.

## Results

### TNF signaling through TNFR1 restricts *C. burnetii* replication within murine macrophages

Our previous studies demonstrated that paracrine TNF produced in response to TLR signaling is critical for restricting *C. burnetii* replication within murine macrophages (27). Consistent with our prior findings, *Tnf^-/-^*BMDMs were significantly more permissive for intracellular bacterial replication than WT BMDMs (Fig. 1A). We also observed that bacterial replication was inhibited completely in both *Tnf*^-/-^ and WT BMDMs that were treated overnight with recombinant murine TNF (rTNF) prior to infection (Fig. 1A). TNF was able to inhibit *C. burnetii* intracellular replication when added as late as 3 days post-infection (Fig. 1B), and *C. burnetii* was able to replicate within macrophages following the removal of rTNF treatment at 24 hours post-infection (Fig. 1B). Collectively, these data indicate TNF inhibits *C. burnetii* replication within BMDMs, in agreement with our previous findings (27).

**Figure 1.**
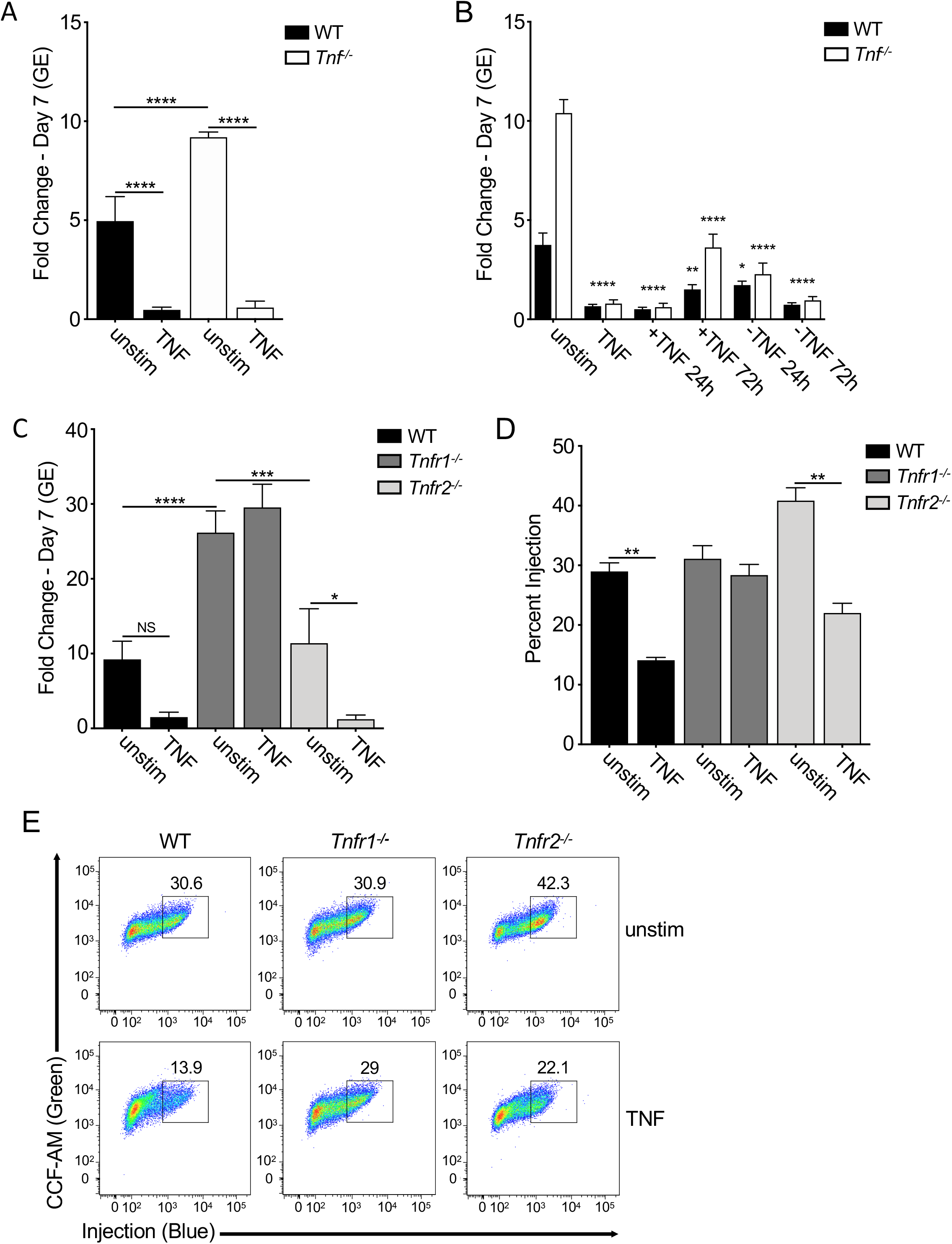
TNF signaling through TNFR1 restricts intracellular *C. burnetii* replication and T4SS effector translocation. (A and B) WT, *Tnf^-/-^*, *Tnfr1^-/-^ and Tnfr2^-/-^* BMDMs were treated overnight with or without 10 ng/ml rTNF and then infected with mCherry-expressing WT *C. burnetii* at an MOI of 50. Bacterial uptake and replication were determined at Day 0 (4hpi) and Day 7, respectively, by measuring genomic equivalents (GE) by qPCR. Bar graphs show fold change relative to Day 0 GE levels ± SEM. Shown are the combined data from four independent experiments, each performed with triplicate wells. (C) WT and *Tnf^-/-^*BMDMs were treated overnight with or without 10 ng/ml rTNF and were then infected with mCherry-expressing WT *C. burnetii* at an MOI of 50. At 24 and 72 hours post-infection, rTNF was add or removed. Bacterial uptake and replication were measured respectively at Day 0 and Day 7 by measuring genomic equivalents (GE) by qPCR. Bar graphs show fold change relative to Day 0 GE levels ± SEM. Each data point compared to fold change growth in unstimulated B6 or *Tnf^-/-^* BMDMs. Shown are the combined data from six independent experiments performed with duplicate or triplicate wells. (D & E) WT, *Tnf^-/-^*, *Tnfr1^-/-^ and Tnfr2^-/-^* BMDMs were treated overnight with or without 10 ng/ml rTNF and then infected with BlaM-CBU_0077-expressing WT or *icmL:Tn C. burnetii* at an MOI of 500. At 24 hours post-infection, BMDMs were loaded with CCF4-AM and analyzed by flow cytometry. (D) Bar graph and (E) flow cytometric plots depicting percent injection are representative of five independent experiments. * = p<0.05, ** = p<0.01, *** = p<0.001, **** = p<0.0001, and NS = no significance.

TNF signals through two distinct receptors, TNF receptor 1 (TNFR1) or TNF receptor 2 (TNFR2). TNFR1 signaling is associated with many potential antimicrobial effector mechanisms, such as inflammatory signaling, induction of cell death, and production of reactive oxygen and nitrogen species, while TNFR2 signaling is associated with cell proliferation, adhesion, and survival (29, 41). Notably, *Tnfr1^-/-^* BMDMs were significantly more permissive for *C. burnetii* intracellular replication than WT BMDMs (Fig. 1C). Moreover, exogenous rTNF addition could not rescue restriction of *C. burnetii* in *Tnfr1^-/-^* BMDMs (Fig. 1C). In contrast, *C. burnetii* replicated within *Tnfr2^-/-^*BMDMs to a similar extent as within WT BMDMs and was similarly restricted by exogenous rTNF treatment. Collectively, these data indicate that TNF signaling through TNFR1, not TNFR2, mediates restriction of *C. burnetii* replication within murine macrophages.

### TNF signaling suppresses T4SS effector translocation

T4SS-dependent translocation of effectors into host cells is necessary for establishment of the *Coxiella-*containing vacuole and subsequent bacterial replication (20). TNF signaling could potentially restrict *C. burnetii* replication by blocking T4SS-dependent translocation of effector proteins. To test the possibility that TNF suppresses T4SS-dependent translocation, we employed a fluorescence resonance energy transfer (FRET)-based reporter system in conjunction with *C. burnetii* harboring a plasmid encoding E. coli TEM-1 ß-lactamase (BlaM) translationally fused to the effector Cbu_0077 (pBlaM-0077) (20). After 24 hours of infection with this strain, BMDMs are loaded with the cell-permeable fluorescent substrate CCF4-AM, which consists of coumarin joined to fluorescein by a β-lactam ring. 409-nm excitation of CCF4-AM results in FRET between coumarin and fluorescein and green fluorescence emission at 518 nm. T4SS-injected BlaM-0077 cleaves CCF4AM and eliminate FRET, resulting in blue fluorescence emission at 447 nm. Injected cells can be distinguished from uninjected cells using flow cytometry.

A similar percentage of WT, *Tnfr1^-/-^*, and *Tnfr2^-/-^* BMDMs were T4SS-injected with BlaM-0077 at 24 hours post-infection (Fig. 1D & E). Notably, rTNF treatment significantly reduced the percentage of WT and *Tnfr2^-/-^* BMDMs that were T4SS-injected compared to unstimulated BMDMs (Figure 1D). In contrast, the percentage of T4SS-injected *Tnfr1^-/-^* BMDMs treated with rTNF was similar to unstimulated BMDMs (Fig. 1D). As expected, BMDMs infected with the *C. burnetii* T4SS-deficient *icmL* mutant carrying pBlaM-0077 were not T4SS-injected (Fig. S1). Taken together, these results indicate that TNF signaling through TNFR1 interferes with T4SS effector translocation and subsequent intracellular bacterial replication.

### TNF-mediated restriction of *C. burnetii* replication occurs independently of cell death and reactive nitrogen and oxygen species

TNF signaling can elicit various antimicrobial programs and cell fates. One such response is the activation of different cell death pathways, which can result in elimination of infected cells (31, 42). TNFR1 signaling can also promote activation of RIPK1’s kinase activity (43), leading to either caspase-8-dependent apoptosis or RIPK3-dependent programmed necrosis (44, 45). To examine whether these death pathways contribute to TNF-dependent restriction of *C. burnetii* infection, we derived BMDMs from *Ripk3^-/-^* or *Casp8^-/-^Ripk3^-/-^*mice. BMDMs from *Casp8^-/-^Ripk3^-/-^* mice must be utilized to examine the role of caspase-8 because *Casp8^-/-^* mice are embryonic lethal (46). Unexpectedly, in untreated BMDMs, *Ripk3^-/-^*BMDMs were significantly more permissive for *C. burnetii* replication, indicating a possible role for RIPK3 in restricting *C. burnetii*. However, exogenous TNF was still able to restrict intracellular *C. burnetii* replication in the absence of RIPK3 or both CASP8 and RIPK3 (Fig. 2A), indicating that both CASP8 and RIPK3 are dispensable for TNF-mediated restriction. BMDMs derived from RIPK1-kinase dead (*Ripk1^KD^*) mice, which lack RIPK1 kinase activity but retain other RIPK1 signaling functions (47), also restricted *C. burnetii* replication following rTNF treatment (Fig. 2B). Following infection with *C. burnetii* expressing BlaM-0077, we observed that the percentage of T4SS-injected cells was also significantly decreased in *Casp8^-/-^Ripk3^-/-^*cells treated with rTNF, similarly to WT BMDMs (Fig. S2). Collectively, these data indicate that the caspase-8-mediated apoptosis and necroptosis pathways do not contribute to TNF-driven restriction of *C. burnetii* T4SS translocation or replication within macrophages.

**Figure 2.**
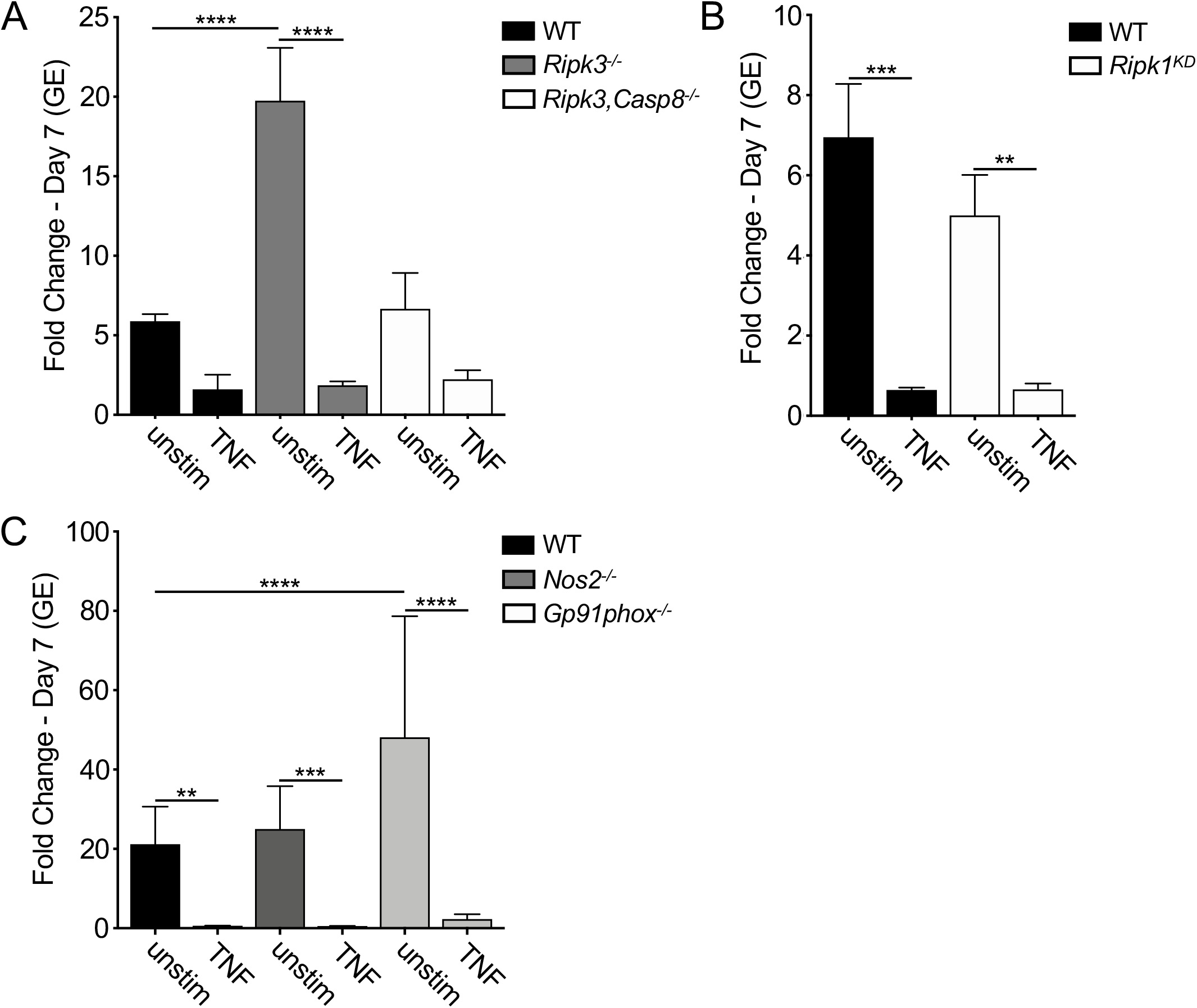
TNF restricts *C. burnetii* replication in BMDMs independent of cell death and ROS/RNS production. (A) WT, *Ripk3^-/-^*, and *Ripk3/Caspase8^-/-^* or (B) WT and *Ripk1^kd/kd^* BMDMs were treated overnight with or without 10 ng/ml rTNF and were then infected with mCherry-expressing WT *C. burnetii* at an MOI of 50. (C) C57BL/6J (WT), *Nos2^-/-^* and *Gp91phox^-/-^* BMDMs were treated overnight with or without 10 ng/ml rTNF and were then infected with mCherry-expressing WT *C. burnetii* at an MOI of 10. Bacterial uptake and replication were measured at Day 0 (4hpi) and Day 7, respectively, by measuring genomic equivalents (GE) by qPCR. Bar graphs show fold change relative to Day 0 GE levels ± SEM. Shown are the combined data from 3 independent experiments with duplicate or triplicate wells. * = p<0.05, ** = p<0.01, *** = p<0.001, **** = p<0.0001, and NS = no significance.

TNF signaling can upregulate expression of inducible nitric oxide synthase (iNOS, also known as NOS2) (48) to generate the reactive metabolite nitric oxide (49), which can restrict *C. burnetii* replication in cell lines (50, 51). TNF can also activate NADPH oxidase to generate superoxide (O_2_-) that can be converted into membrane-permeable hydrogen peroxide (H_2_O_2_) (52), which can restrict the growth of various intracellular bacteria, including *C. burnetii* (50, 53). To determine whether NADPH oxidase and/or iNOS play a role in TNF-mediated restriction of *C. burnetii*, we infected BMDMs from mice lacking *Nos2* or *Gp91phox,* which encodes a subunit of NADPH oxidase (52). Though *Gp91phox^-/-^* macrophages were generally more permissive than WT BMDMs, both *Gp91phox^-/-^* and *Nos2^-/-^* BMDMs still restricted *C. burnetii* intracellular replication to levels similar to those observed in WT BMDMs following rTNF treatment (Fig. 2C). Consistently, TNF treatment also restricted *C. burnetii* T4SS injection in *Gp91phox^-/-^* and *Nos2^-/-^* BMDMs comparably to WT BMDMs (Fig. S2). These data indicate that neither iNOS nor NADPH oxidase is required for TNF-driven restriction of *C. burnetii* T4SS translocation or replication.

### Type I interferon and TNF collaborate to restrict *C. burnetii* intracellular replication

We have previously found that *C. burnetii* infection of BMDMs induces low, but biologically active levels of type I IFN (27). Like TNF, type I IFNs can induce antimicrobial programs in host cells to restrict bacterial replication and spread (54). Typically, type I IFNs are induced downstream of microbial stimulation of TLRs or cytosolic sensors (55, 56). Furthermore, type I IFNs can synergize with TNF in both a paracrine and autocrine manner to strongly induce and maintain immune responses during infection (57). To investigate the potential role of type I IFNs in promoting TNF-mediated restriction of *C. burnetii* infection, we infected BMDMs lacking the type I IFN receptor IFNAR1. We found that while *Ifnar^-/-^* BMDMs were significantly more permissive than WT BMDMs, intracellular *C. burnetii* replication was still significantly reduced by TNF treatment (Fig. 3A & B). This restriction, however, was not to the same extent observed with WT BMDMs, as *C. burnetii* replicated within TNF-treated *Ifnar^-/-^* BMDMs to the same extent as untreated WT BMDMs (Fig. 3A). Interestingly, *Coxiella* T4SS injection levels were significantly higher in *Ifnar^-/-^* BMDMs compares to WT, yet both WT and *Ifnar^-/-^*BMDMs showed similar decreases in the percentage of T4SS-injected cells following TNF treatment (Fig. 3C & D), indicating that TNF restricts T4SS translocation independently of IFNAR1 signaling.

**FIgure 3.**
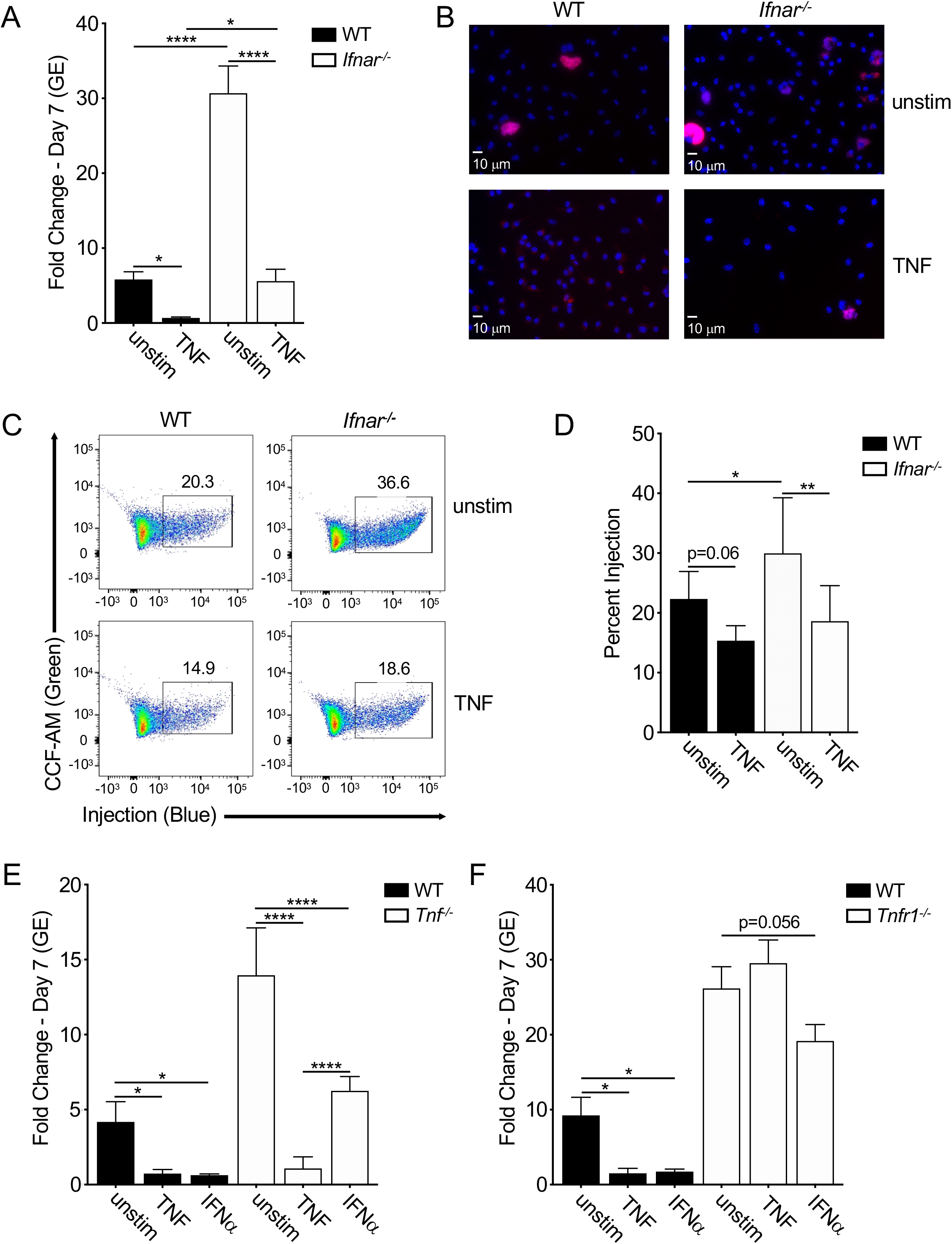
TNF works in concert with type I IFN to restrict *C. burnetii* T4SS translocation and replication in murine macrophages. (A) WT and *Ifnar^-/-^* BMDMs were treated overnight with or without 10 ng/ml rTNF and were then infected with mCherry-expressing WT *C. burnetii* at an MOI of 50. Bacterial uptake and replication were determined at Day 0 (4hpi) and Day 7, respectively, by measuring genomic equivalents (GE) by qPCR. Bar graphs show fold change relative to Day 0 GE levels ± SEM. Shown are the combined data from ten independent experiment with duplicate or triplicate wells. (B) Representative fluorescence micrographs of infected BMDMs stained with DAPI on Day 7 post-infection at 40X magnification. (C & D) WT and *Ifnar^-/-^* BMDMs were treated overnight with and without 10 ng/ml rTNF and were then infected with BlaM-CBU_0077-expressing WT or icmL:Tn *C. burnetii* at an MOI of 500. At 24 hours post-infection, BMDMs were loaded with CCF4-AM and analyzed through flow cytometry. Shown are (C) flow cytometric plots representative of three independent experiments and (D) bar graph of combined data from three independent experiments. (E) C57BL/6 and *Tnf^-/-^*BMDMs or (F) WT and *Tnfr1^-/-^* BMDMs were treated overnight with or without 10 ng/ml rTNF or 10^3^ U/ml rIFNα and were then infected with mCherry-expressing WT *C. burnetii* at an MOI of 50. Bacterial uptake and replication were measured at Day 0 (4hpi) and Day 7, respectively, by measuring genomic equivalents (GE) by qPCR. Bar graphs show fold change relative to Day 0 GE levels ± SEM. Data from four independent experiment with duplicate or triplicate wells. * = p<0.05, ** = p<0.01, *** = p<0.001, **** = p<0.0001, and NS = no significance.

To further define the effects of type I IFN on *C. burnetii* replication, we treated WT BMDMs with recombinant interferon-α (rIFNα) and observed suppression of bacterial restriction similar to that observed with TNF treatment (Fig. 3E). Interestingly, rIFNα treatment of *Tnf^-/-^* BMDMs led only to partial restriction of *C. burnetii* replication (Fig. 3E), indicating that TNF is required for maximal restriction of *C. burnetii* by type I IFN. We similarly observed that *Tnfr1^-/-^* BMDMs treated with rIFNα were unable to fully restrict *C. burnetii* replication (Fig. 3F). Taken together, these data indicate that type I IFN synergizes with TNF signaling to restrict *C. burnetii* replication within macrophages.

### TNF and IFN upregulate IRG1 to suppress *C. burnetii* replication in murine macrophages

Given that both TNF and type I IFN are required to restrict *C. burnetii* within murine macrophages, we next considered antimicrobial host factors that could be regulated by both TNF and type I IFN. The enzyme immune-responsive gene 1 (IRG1) also known as cis-aconitate decarboxylase 1 (ACOD1) catalyzes the conversion of the TCA cycle intermediate aconitate into itaconate and is upregulated by a variety of pro-inflammatory cytokines, including both TNF and type I IFN (35, 58). IRG1 contributes to the restriction of intracellular pathogens, including *Salmonella enterica*, *M. tuberculosis* and *L. pneumophila* (59, 60). In both WT and *Tnf^- /-^* BMDMs, *Irg1* mRNA expression was induced by either exogenous TNF treatment or *C. burnetii* infection, and it was induced even more so by both together (Fig. 4A). *Irg1* was also induced in *Ifnar^-/-^* BMDMs following TNF stimulation and *C. burnetii*, but to a lower extent than in WT BMDMs (Fig. 4A), indicating that endogenous type I IFN signaling is required for maximal TNF-dependent Irg1 expression. *Irg1* induction was also further increased by combined rIFNα and *C. burnetii* infection in both WT and *Tnf^-/-^* BMDMs, but not in *Ifnar^-/-^* BMDMs (Fig. 4A). These data indicate that *C. burnetii* infection induces *Irg1* expression, and that Irg1 expression is further enhanced by TNF and type I IFN stimulation.

**Figure 4.**
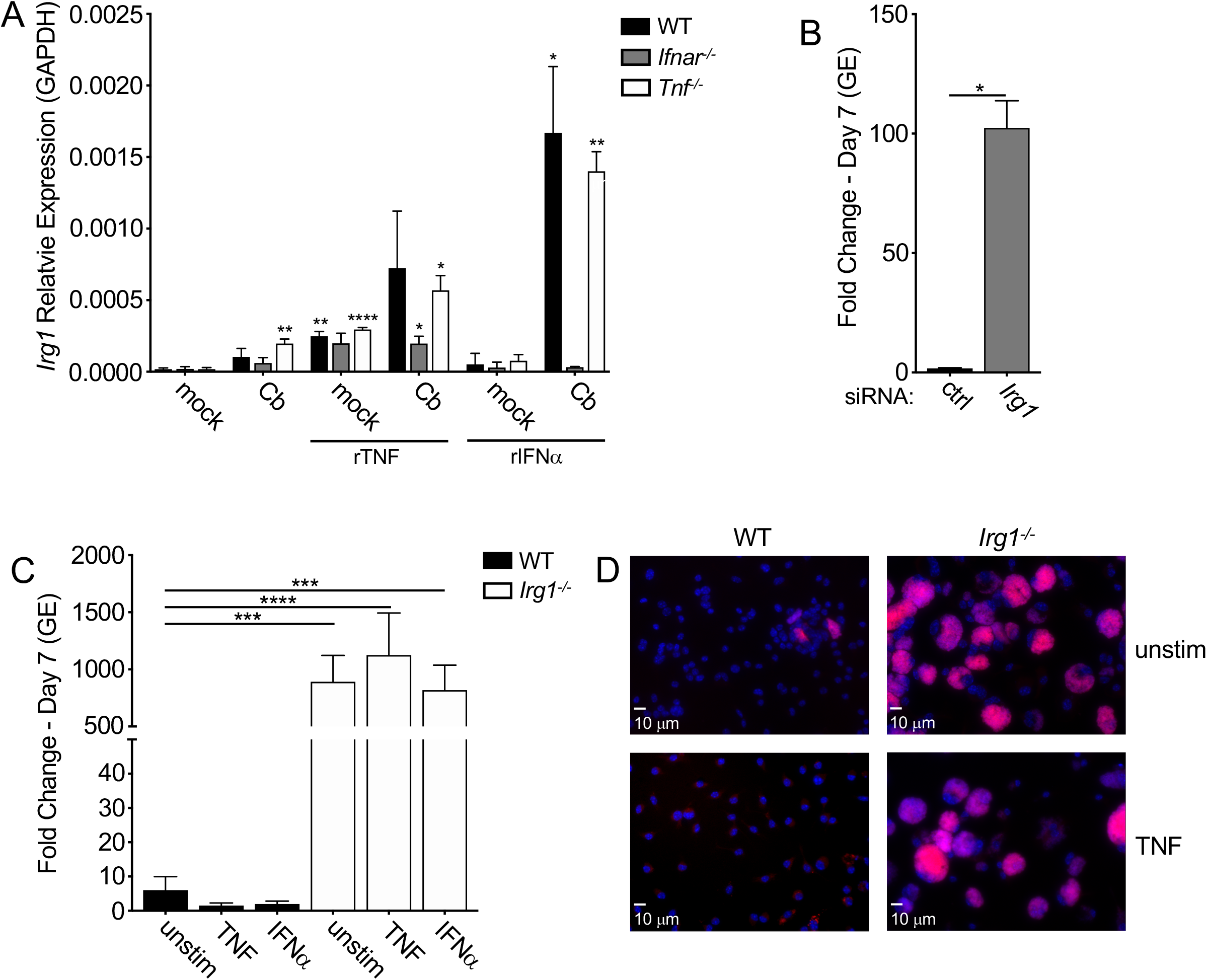
TNF and IFNα act through IRG1 to suppress *C. burnetii* replication in BMDMs. (A) WT, *Tnf^-/-^* and *Ifnar^-/-^* BMDMs were treated with or without 10 ng/ml rTNF or 10^3^ U/ml rIFNα and were then mock-infected or infected with mCherry-expressing WT *C. burnetii* at an MOI of 50. Expression of *Irg1* mRNA relative to the housekeeping gene *Gapdh* was measured by qPCR. Bar graphs show mean relative expression ± SEM from triplicate wells. Representative of two independent experiments. Statistical significance based on comparison to relative expression in mock untreated cells. (B) WT BMDMs were stimulated with 10 ng/ml rTNF and transfected with control scrambled or *Irg1*-targeting siRNA prior to infection with mCherry-expressing WT *C. burnetii* at an MOI of 50. Bacterial uptake and replication were measured at Day 0 and Day 7, respectively, by measuring genomic equivalents (GE) by qPCR. Bar graphs show fold change relative to Day 0 GE levels ± SEM. Shown are the combined data of 4 independent experiments with triplicate wells. (C) WT and *Irg1^-/-^* BMDMs were treated overnight with or without 10 ng/ml rTNF or 10^3^ U/ml rIFNα and then infected with mCherry-expressing WT *C. burnetii* at an MOI of 50. Bacterial uptake and replication were measured at Day 0 and Day 7, respectively, by measuring genomic equivalents (GE) by qPCR. Bar graphs show fold change relative to Day 0 GE levels ± SEM. Shown are the combined data from 4 independent experiments with duplicate or triplicate wells. (D) Representative fluorescence micrographs of infected BMDMs were fixed and stained with DAPI on Day 7 post-infection and imaged at 40X magnification. * = p<0.05, ** = p<0.01, *** = p<0.001, **** = p<0.0001, and NS = no significance.

To test the possible contribution of IRG1 to restricting *C. burnetii* intracellular replication, we transfected WT BMDMs with siRNA targeting *Irg1* or a scrambled control siRNA prior to and during *C. burnetii* infection. Notably, *Irg1* siRNA treatment resulted in a 70-80% knockdown of *Irg1* mRNA levels over the course of a one-week infection (Fig. S3A). Critically, *Irg1* knockdown resulted in a significant increase in *C. burnetii* intracellular replication compared to control siRNA-treated cells (Fig. 4B, S3B). Furthermore, *Irg1^-/-^* BMDMs exhibited a significant increase in *C. burnetii* replication compared to WT BMDMs (Fig. 4C and D). Importantly, treatment of *Irg1^-/-^* BMDMs with either TNF (Fig. 4C and D) or IFNα (Fig. 4C) failed to restrict *C. burnetii* replication. Taken together, these results indicate that IRG1 is required for TNF- and type I IFN-driven restriction of *C. burnetii* intracellular replication in murine macrophages.

### Itaconic acid restricts *C. burnetii* intracellular replication

*IRG1* encodes a cis-aconitate decarboxylase (ACOD1) that converts the TCA cycle metabolite aconitate into itaconate, also known as itaconic acid (ITA). ITA inhibits the metabolism and growth of bacterial pathogens by targeting critical bacterial metabolic enzymes, including isocitrate lysase, part of the glyoxylate shunt pathway, and methylisocitrate lyase in the 2-methylcitrate cycle (59-62). To test whether exogenous ITA can complement IRG1 deficiency and restore restriction of *C. burnetii* intracellular replication in *Irg1^-/-^* BMDMs, we treated WT or *Irg1^-/-^*BMDMs with low (1 mM) as well as high (10 mM) doses of ITA. Importantly, 10 mM ITA is compatible with physiological doses, as it is similar to the concentration of ITA found in activated RAW264.7 murine macrophage cells (59). Critically, ITA treatment potently restricted *C. burnetii* replication within both WT and *Irg1^-/-^* BMDMs in a dose-dependent manner (Fig. 5A and B). We also found that high dose ITA inhibited *C. burnetii* replication even when added as late as 72 hours post-infection (Figs. 5C, S4). We observed that removal of 1mM, but not 10mM, ITA treatment at 24 or 72 hours following *C. burnetii* infection allowed for some recovery of bacteria replication, suggesting that ITA acts in a bacteriostatic manner to limit replication (Fig. 5C and S4). Altogether, these data indicate that ITA restricts *C. burnetii* replication within macrophages.

**Figure 5.**
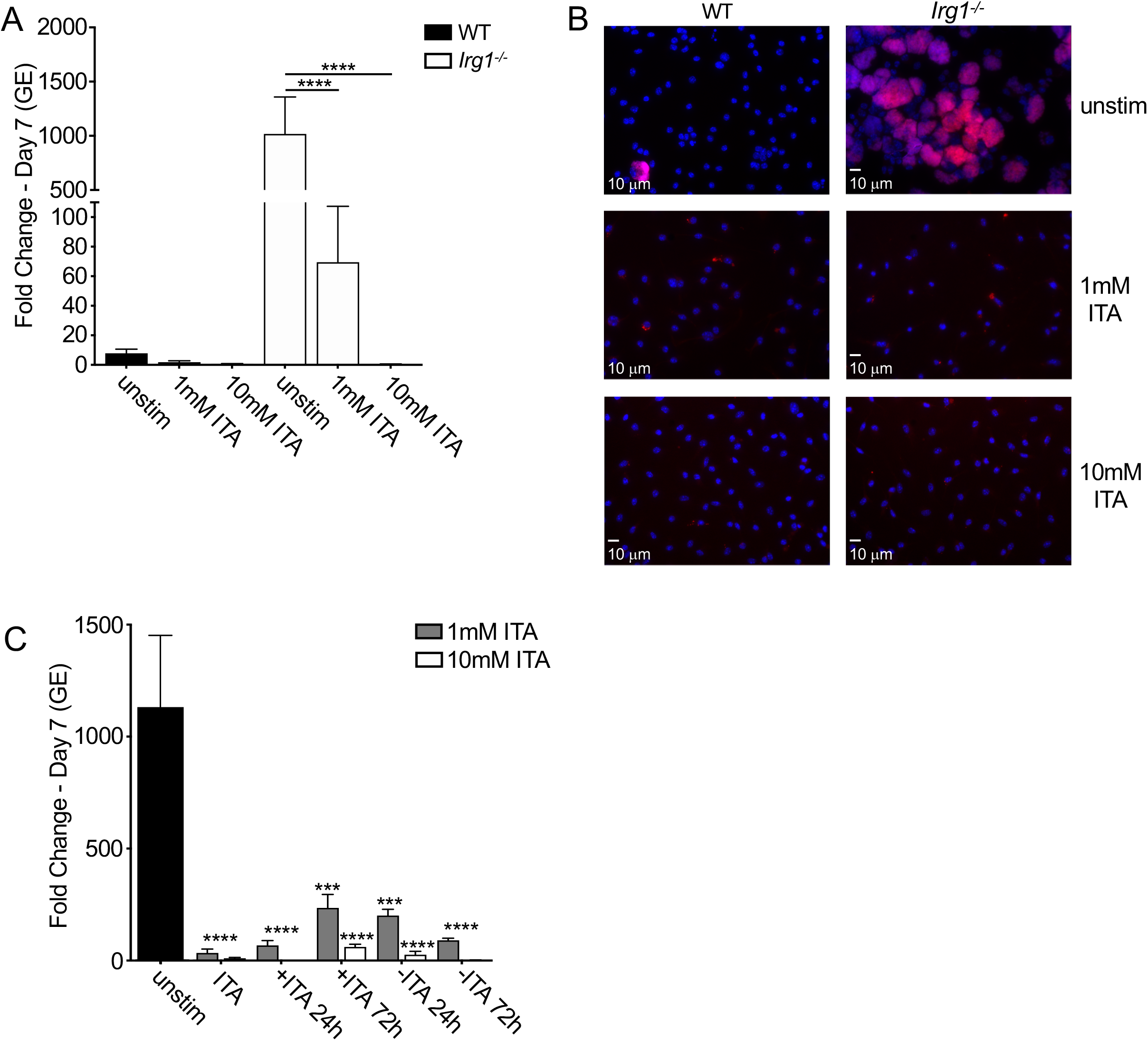
Itaconic acid restricts *C. burnetii* intracellular replication in mouse macrophages. WT and *Irg1^-/-^* BMDMs were treated with and without 1mM or 10mM itaconic acid (ITA) and were then infected with mCherry-expressing WT *C. burnetii* at an MOI of 50. For (C), ITA was added or removed at 24 and 72 hours post-infection in *Irg1^-/-^*BMDMs. (A and C) Bacterial uptake and replication were measured at Day 0 and Day 7, respectively, by measuring genomic equivalents (GE) by qPCR. Bar graphs show fold change relative to Day 0 GE levels ± SEM. Shown are the combined data from (A) 5 or (C) 3 independent experiments with duplicate or triplicate wells. (B) Representative fluorescence micrographs of infected BMDMs were stained with DAPI on Day 7 post-infection and imaged at 40X magnification. * = p<0.05, ** = p<0.01, *** = p<0.001, **** = p<0.0001, and NS = no significance.

### Itaconic acid inhibits *C. burnetii* T4SS translocation and axenic replication

Given that TNF and IFN treatment limit *C. burnetii* T4SS translocation, we considered the possibility that they do so via production of IRG1-dependent ITA. Indeed, a significantly higher percentage of *Irg1^-/-^* BMDMs were injected compared to WT BMDMs at 24 hours following infection with *C. burnetii* expressing BlaM-0077, demonstrating that the absence of IRG1 correlates with increased translocation (Fig. 6A & B). Moreover, IRG1 was required for TNF-dependent blockade of T4SS injection, as TNF treatment was unable to inhibit T4SS injection in *Irg1^-/-^* BMDMs (Fig. 6A). Critically, ITA treatment of either WT or *Irg1^-/-^* BMDMs during infection with BlaM-0077-expressing *C. burnetii* significantly reduced T4SS injection in both genotypes (Fig. 6A). Collectively these data indicate that ITA and IRG1 mediate TNF-induced blockade of *C. burnetii* T4SS-injection within BMDMs.

**Figure 6.**
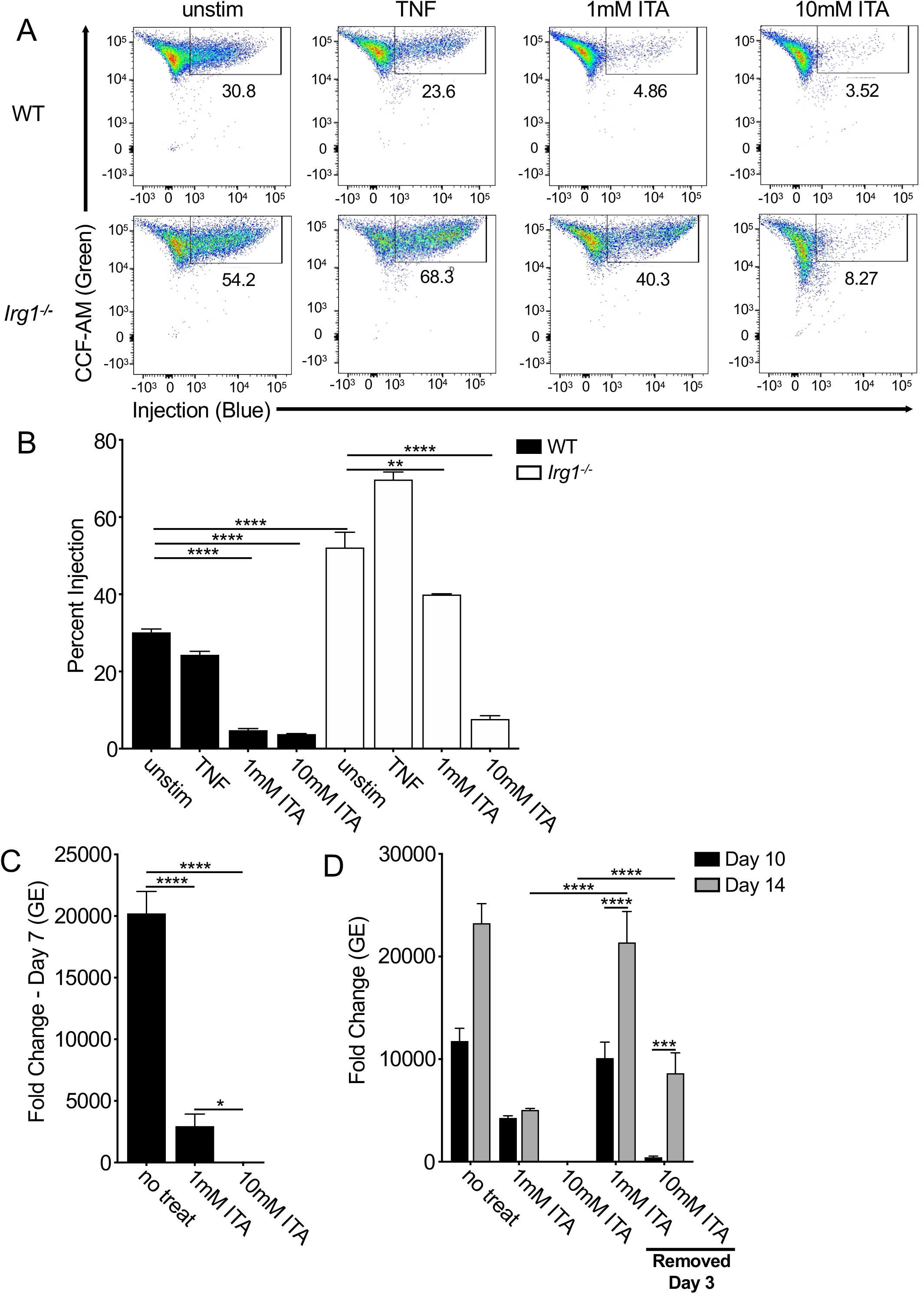
Itaconic acid inhibits *C. burnetii* T4SS injection and axenic replication. (A & B) WT and *Irg1^-/-^* BMDMs, with or without 10 ng/ml rTNF or 1mM or 10mM itaconic acid (ITA), were infected with BlaM-CBU_0077-expressing WT or icmL:Tn *C. burnetii* at an MOI of 500. At 24 hours post-infection, BMDMs were loaded with CCF4-AM and analyzed through flow cytometry. Shown are (A) representative flow plots and (B) bar graphs depicting the combined data from three independent experiments. (C and D) mCherry-expressing WT *C. burnetii* was inoculated into ACCM-2 media, with or without ITA, and incubated for 7 days; for inoculate with ITA, pH was readjusted prior to inoculation with NaOH. In (D), ITA was removed at Day 3 for some inoculates by centrifugation, washing and resuspension in new ACCM-2. At Day 7, 10 and 14, replication was measured by GE via qPCR. Bar graphs show fold change relative to starting inoculate concentration of 2 × 10^6^ GE/ml. Shown are the combined data from four independent experiment. * = p<0.05, ** = p<0.01, *** = p<0.001, **** = p<0.0001, and NS = no significance.

**Figure 7.**
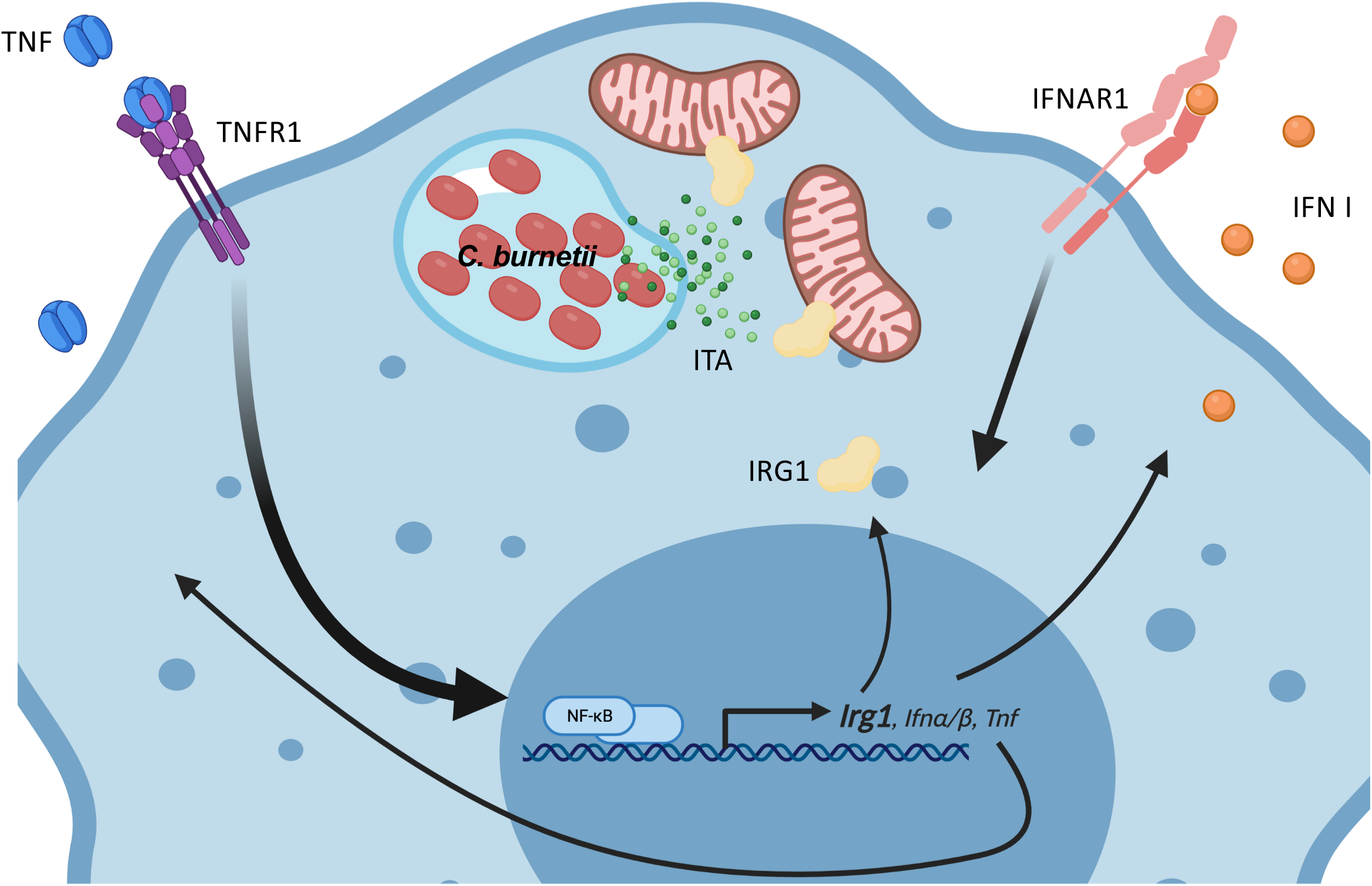
Graphical model of TNF and type I IFN signaling upregulating IRG1, which produces itaconic acid to restrict *C. burnetii* replication in mouse macrophages. Infection of macrophages by *C. burnetii* elicits TLR-driven production of TNF, which is secreted extracellularly and signals in a paracrine and autocrine manner. TNF signaling through TNFR1 and type I IFN signaling through IFNAR work in concert to upregulate expression of IRG1, which localizes to the mitochondria to catabolize production of itaconic acid (ITA), which subsequently blocks the replication of *C. burnetii* in the host cell. Graphical model created with BioRender.com.

ITA could inhibit *Coxiella* replication by acting directly on the bacteria, or by enhancing host immune responses. Given that ITA inhibits bacterial metabolic enzymes in multiple bacterial species (59-62), we tested the direct effect of ITA on *C. burnetii* growth outside of the host cell. We observed that ITA inhibited *C. burnetii* axenic replication almost completely at concentrations of 1 and 10 mM (Fig. 6C). Interestingly, *C. burnetii* grown for 3 days in media containing 1mM ITA fully recovered and replicated to high levels when moved to ITA-free media (Fig. 6C); *C. burnetii* was also able to recover and replicate to some extent after 3 days of incubation in media containing 10mM ITA (Fig. 6D). Taken together, these results indicate that ITA has a direct, dose-dependent inhibitory effect on *C. burnetii* axenic replication.

## Discussion

Tumor necrosis factor (TNF) signaling is a crucial arm of the antimicrobial immune response that mediates restriction of various intracellular bacterial pathogens, including *M. tuberculosis*, *L. monocytogenes, L. pneumophila,* and *C. burnetii*. We show here that TNF synergizes with type I IFNs for maximal restriction of *C. burnetii* through induction of IRG1. We also found that the IRG1-produced metabolite itaconate (ITA) restricted *C. burnetii* T4SS translocation and replication within macrophages and during axenic growth. Thus, TNF and type I IFN function through the IRG1-itaconate pathway to restrict *C. burnetii* replication within the macrophages.

We found that TNF and type I IFN signaling work together to restrict *C. burnetii* within BMDMs. BMDMs deficient in either TNF or type I IFN signaling were more permissive for *C. burnetii* replication, indicating that endogenous production of TNF and type I IFN in response to *C. burnetii* mediates this restriction. Indeed, we have previously found that *C. burnetii* infection of BMDMs induced very low, albeit biologically active, levels of type I IFN (27). Interestingly, type I IFN signaling also restricts *C. burnetii* during *in vivo* infection of mice (63). We also found that exogenous type I IFN was able to restrict *C. burnetii* in BMDMs as well as exogenous TNF in wild-type cells, but type I IFN and TNF were reciprocally dependent on each other for full restriction. The synergistic relationship of TNF and type I IFN has been shown before in the context of other pathogens like vesicular stomatitis virus (VSV) and Sendai virus (64, 65), and type I IFNs themselves are crucial for resistance against the bacterial pathogens group B streptococci (GBS), *S. pneumoniae* and *E. coli,* resulting from defective TNF and IFNγ in the absence of IFN signaling (66). Notably, TNF and type I interferons can operate in an autocrine loop, whereby TNF signaling stimulates the production and release of IFNs, which can then act back upon the cell and reinforce the immune response (57). Perhaps this type of autocrine loop also contributes to the ability of TNF and type I IFN to restrict *C. burnetii* replication via upregulation of Immune Responsive Gene 1 (*Irg1*).

We also found that TNF and type I IFN signaling upregulate expression of the enzyme IRG1 (Immune-Responsive Gene 1). Furthermore, IRG1 was necessary for TNF and type I IFN-mediated restriction of *C. burnetii* intracellular replication in murine bone marrow-derived macrophages. IRG1 has emerged as a key immune effector against both viral and bacterial pathogens, including Zika virus, *S. enterica*, *M. tuberculosis*, *L. pneumophila* (59, 60, 67). IRG1 converts the TCA metabolite aconitate into itaconate (itaconic acid or ITA). ITA is one of the most abundant metabolites found in activated macrophages and possesses competitive inhibitory properties against various enzymes. Critically, we found that exogenous ITA potently inhibited *C. burnetii* replication within macrophages at biologically relevant doses (59). Furthermore, ITA potently restricted *C. burnetii* axenic growth, indicating a direct inhibitory effect by ITA on *C. burnetii*. Our findings with IRG1 and ITA restricting *C. burnetii* replication within murine macrophages are in agreement with recent findings by another group (68). They found that *C. burnetii* infection induced IRG1 expression in a MyD88-dependent manner, and that IRG1 and ITA restrict *C. burnetii* replication within murine macrophages. Our data indicate that TNF is the likely candidate cytokine produced downstream of TLR/MyD88 signaling that drives IRG1 expression. Furthermore, they also found that *Irg1^-/-^*mice have a defect in control of *C. burnetii* infection *in vivo* and treatment of *Irg1^-/-^*mice with ITA reduced *C. burnetii* bacterial loads (68). Interestingly, they found that human macrophages produce much less itaconate than murine macrophages, which may explain why human macrophages are much more permissive for *C. burnetii* replication than murine macrophages (68).

ITA has direct antibacterial effects on other bacterial species, at least in part by inhibiting bacterial metabolic enzymes such as isocitrate lysase, which is part of the glyoxylate shunt pathway, propionyl-CoA carboxylase in the citramalate cycle, as well as methylisocitrate lyase in the 2-methylcitrate cycle (59, 61, 62, 69-71). Interestingly, like its close evolutionary relative *L. pneumophila*, *C. burnetii* lacks a glyoxylate shunt, but does encode a methylisocitrate lyase (*prpB*). In the future, it would be of interest to determine whether *prpB* or other bacterial enzymes are the potential targets of itaconate in *C. burnetii*.

*C. burnetii* possesses a number of effectors that block cell death (72-75), and *C*. *burnetii-*infected BMDMs do not undergo cell death (27). Consistently, TNF-driven restriction of *C. burnetii* is not due to CASP8 or RIPK3-mediated cell death of infected cells. Intriguingly, our recent findings reveal that TNF restriction of the evolutionarily related intracellular pathogen *L. pneumophila* acts through inflammasome-mediated cell death involving caspases-1, 8, and 11 (76). Unlike *C. burnetii,* which possesses several effectors that potently block apoptosis and inflammasome-mediated cell death, *L. pneumophila* has not coevolved with mammalian hosts, and therefore does not have effectors that robustly block regulated cell death in mammalian macrophages. Thus, these findings highlight functional differences in how TNF restricts these two pathogens, possibly due to differences in the how these two pathogens target host pathways.

We also found, surprisingly, that NADPH oxidase (NOX) and inducible nitric oxide synthase (iNos) were dispensable for TNF-mediated restriction of *C. burnetii* in BMDMs. Importantly, however, in the absence of exogenous TNF treatment, *Gp91phox^-/-^* BMDMs were significantly more permissive than WT BMDMs, indicating that while NADPH oxidase does not mediate TNF-induced restriction, NADPH oxidase does contribute to basal control of *C. burnetii*. Notably, *Gp91phox^-/-^* mice are also more susceptible to for *C. burnetii* infection (50), highlighting a greater role for ROS in the context of *C. burnetii* infection. Potentially, these pathways are complementary or redundant; for example, iNOS has recently been shown to act redundantly with five other immune genes, including IRG1, to restrict *L. pneumophila* downstream of IFNγ (60).

In conclusion, our findings indicate that TNF and type I IFN signaling work in concert to upregulate expression of the antimicrobial immunomodulatory enzyme IRG1 to produce the antimicrobial metabolite itaconate, which restricts intracellular *C. burnetii* replication within mouse macrophages. Collectively, our findings provide new insight into the mechanisms by which TNF signaling promotes control of intracellular bacterial pathogens.

## Acknowledgements

We thank Igor Brodsky and Dieter Schifferli and members of the Shin and Brodsky labs for helpful scientific discussions and advice. We thank William Bradley for experimental protocols, Jianxin You for use of her fluorescence microscope, and Xin Liu for technical advice on flow cytometry. This study was funded by National Institutes of Health grants R01AI118861, R01AI123243 (S.S.), and T32GM07229 (N.L.F.), a Burroughs-Wellcome Fund Investigators in the Pathogenesis of Infectious Diseases Award (S.S.), and the HHMI James H. Gilliam, Jr. Fellowship for Advanced Study (N.L.F.).

## FIGURES LEGEND

### Supplemental Figures

**Figure S1. related to Figure 1.**
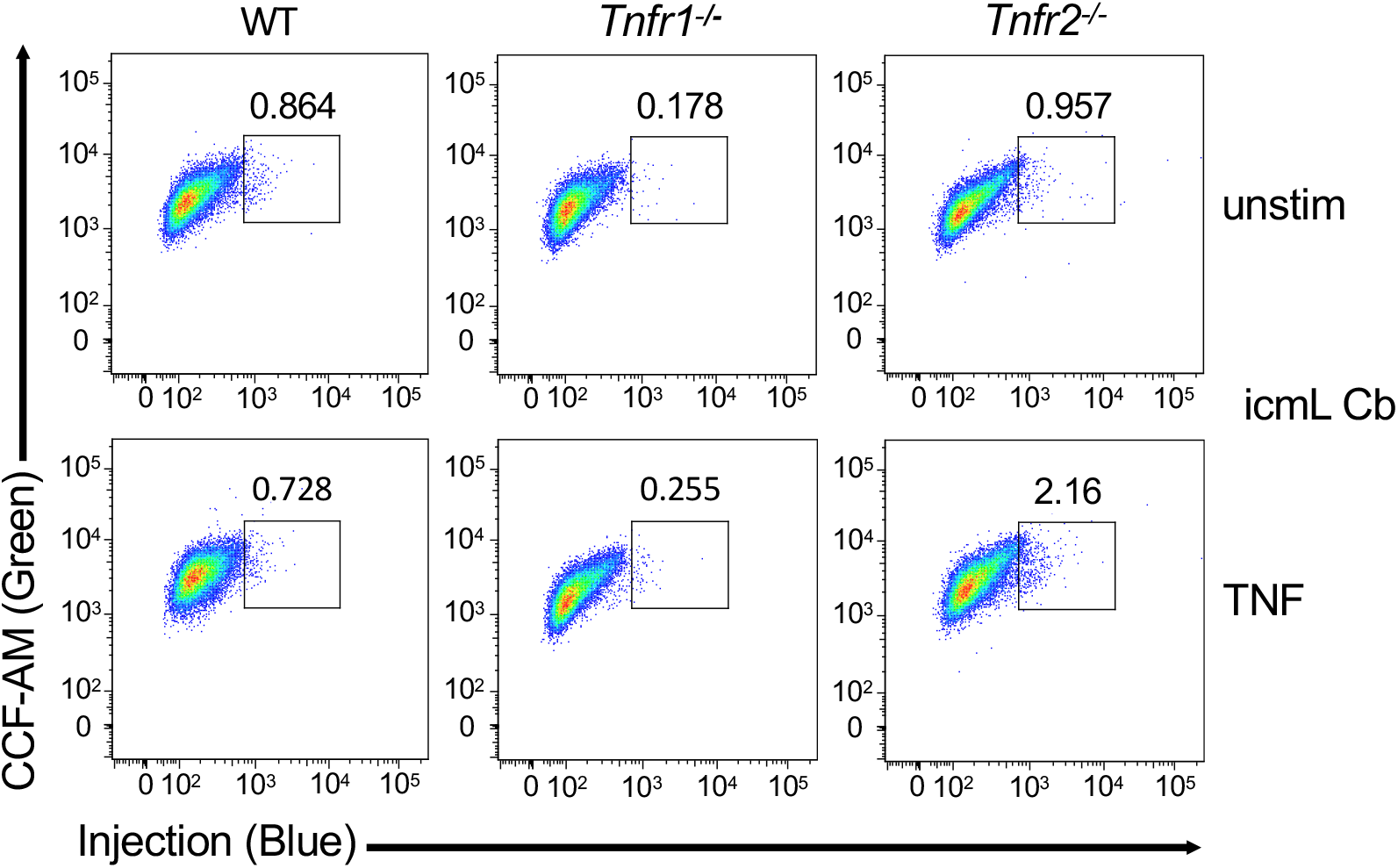
T4SS is required for *C. burnetii* translocation of CBU_0077 into murine BMDMs. C57BL/6J (WT), *Tnfr1^-/-^,* and *Tnfr2^-/-^* BMDMs, with and without 10 ng/ml rTNF, were infected with BlaM-CBU_0077-expressing *icmL:Tn C. burnetii* at an MOI of 500. At 24 hours post-infection, BMDMs were loaded with CCF4-AM and analyzed by flow cytometry. Shown are representative flow cytometric plots. Bar graph depicting percent injection is showing the combined data from five independent experiments.

**Figure S2. related to Figure 2.**
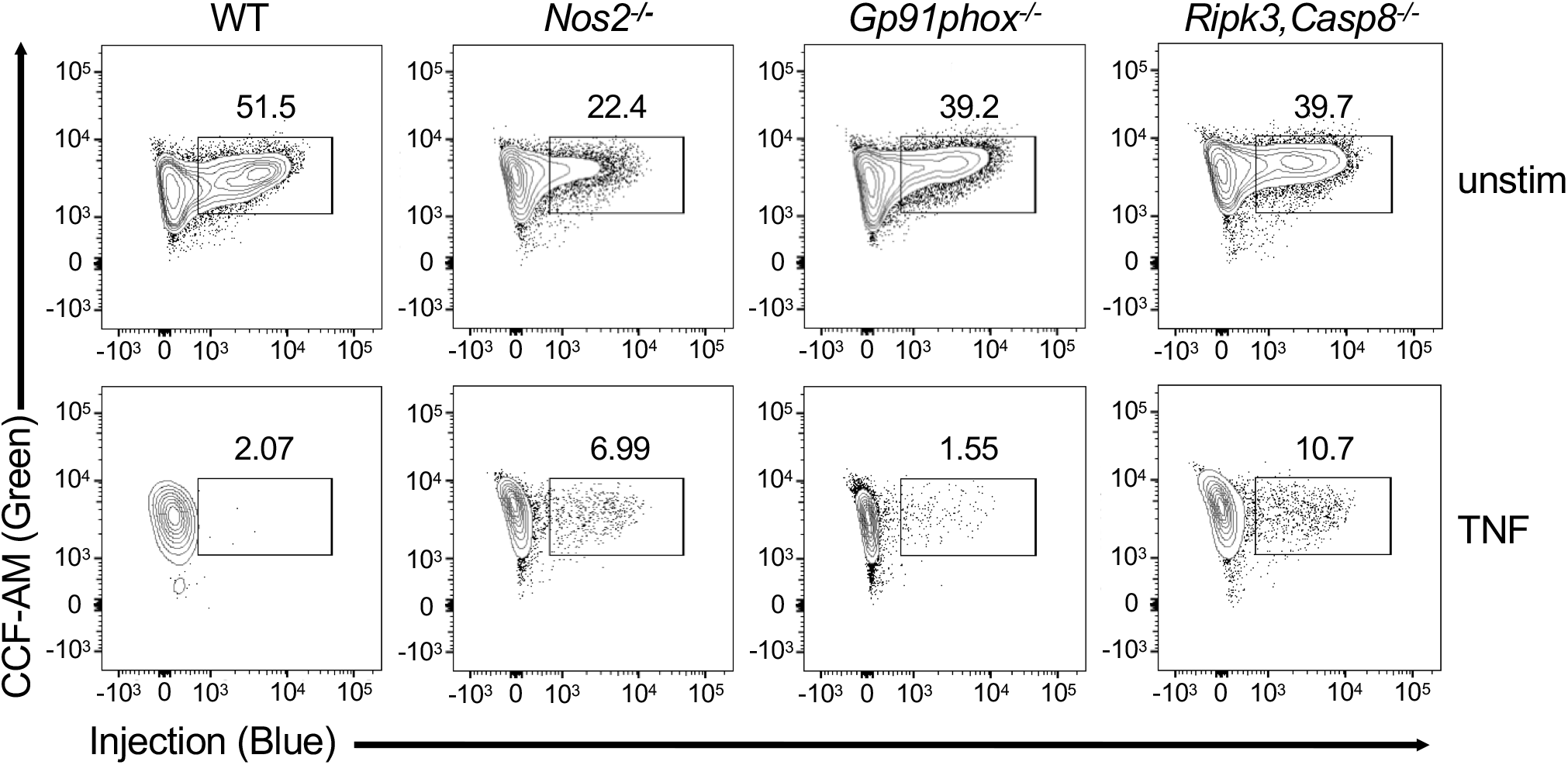
TNF restricts *C. burnetii* T4SS injection in murine macrophages independently of ROS and RNS production and RIPK3- and caspase-8-mediated cell death. C57BL/6J (WT), *Nos2^-/-^, Gp91phox^-/-^,* and *Ripk3^-/-^Casp8^-/-^* BMDMs were infected with BlaM-CBU_0077-expressing WT or *icmL:Tn C. burnetii* at an MOI of 500. At 24 hours post-infection, BMDMs were loaded with CCF4-AM and analyzed by flow cytometry. Flow cytometric plots are representative of three independent experiments.

**Figure S3. related to Figure 4.**
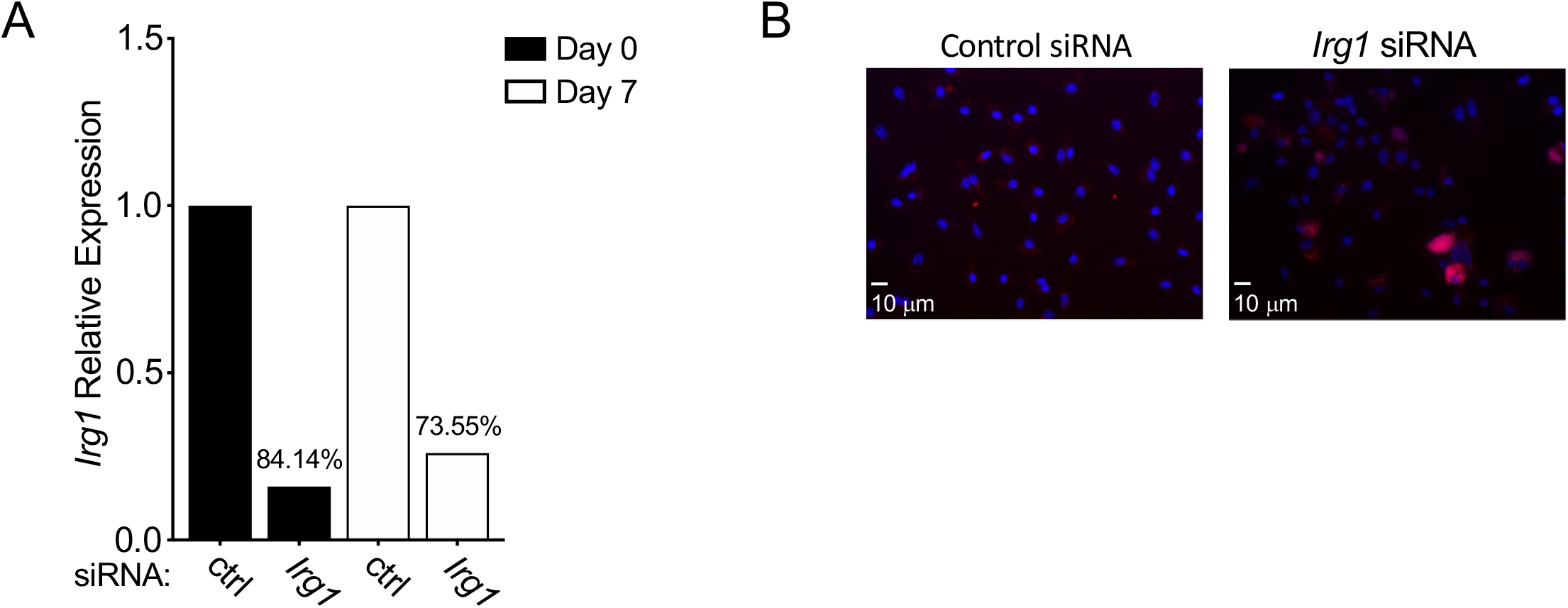
siRNA-mediated silencing of *Irg1* and fluorescent imaging of murine BMDMs infected with *C. burnetii*. (A) C57BL/6 (WT) BMDMs were stimulated overnight with 10 ng/ml rTNF and transfected with control scrambled or *Irg1*-targeting siRNA. They were then infected with mCherry-expressing WT *C. burnetii* at an MOI of 50. *Irg1* knockdown efficiency was analyzed at Day 0 and Day 7 by by qPCR. Bar graphs show *Irg1* mRNA expression relative to the housekeeping gene *Gapdh*. Representative graph of 4 independent experiments. (B) Representative fluorescence micrographs of infected WT BMDMs treated with control scrambled or *Irg1* siRNA treatment. BMDMs were fixed and stained with DAPI on Day 7 post-infection and imaged at 40X magnification.

**Figure S4. related to Figure 5.**
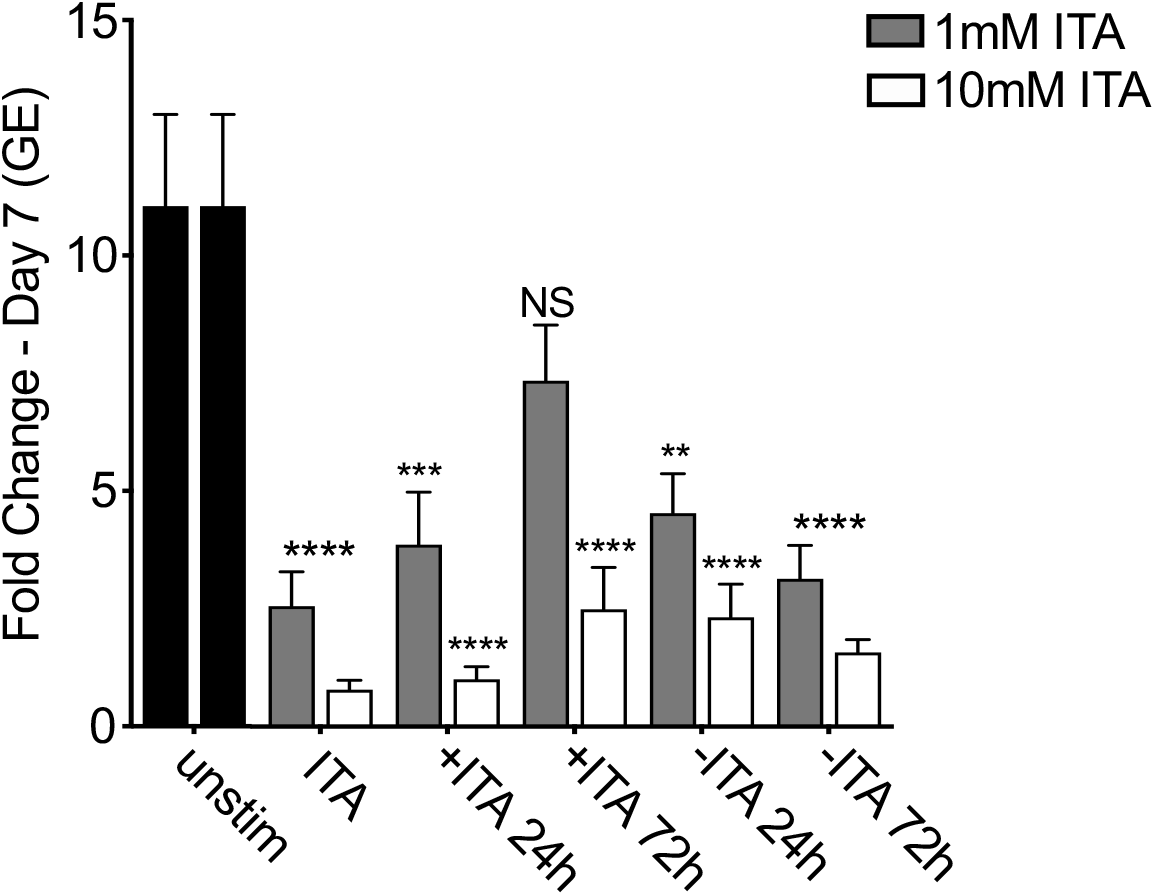
Itaconic acid restricts *C. burnetii* intracellular growth. WT BMDMs were treated with or without 1mM or 10mM itaconic acid (ITA) and were then infected with mCherry-expressing WT *C. burnetii* at an MOI of 50. ITA was added or removed at 24 and 72 hours post-infection. Bacterial uptake and replication were measure respectively at Day 0 and Day 7 by measuring genomic equivalents (GE) by qPCR. Bar graphs show fold change relative to Day 0 GE levels ± SEM. Shown are the combined data from 3 independent experiment with triplicate wells.

## References

1. Tissot-Dupont H, Raoult D. Q fever. Infect Dis Clin North Am. 2008;22(3):505–14, ix. doi: 10.1016/j.idc.2008.03.002. PubMed PMID: 18755387.

2. Carcopino X, Raoult D, Bretelle F, Boubli L, Stein A. Q Fever during pregnancy: a cause of poor fetal and maternal outcome. Ann N Y Acad Sci. 2009;1166:79–89. doi: 10.1111/j.1749-6632.2009.04519.x. PubMed PMID: 19538266.

3. Eldin C, Mélenotte C, Mediannikov O, Ghigo E, Million M, Edouard S, et al. From Q Fever to Coxiella burnetii Infection: a Paradigm Change. Clin Microbiol Rev. 2017;30(1):115–90. doi: 10.1128/CMR.00045-16. PubMed PMID: 27856520; PubMed Central PMCID: PMC5217791.

4. van der Hoek W, Dijkstra F, Schimmer B, Schneeberger PM, Vellema P, Wijkmans C, et al. Q fever in the Netherlands: an update on the epidemiology and control measures. Euro Surveill. 2010;15(12). Epub 2010/03/25. PubMed PMID: 20350500.

5. Vanderburg S, Rubach MP, Halliday JE, Cleaveland S, Reddy EA, Crump JA. Epidemiology of Coxiella burnetii infection in Africa: a OneHealth systematic review. PLoS Negl Trop Dis. 2014;8(4):e2787. Epub 2014/04/10. doi: 10.1371/journal.pntd.0002787. PubMed PMID: 24722554; PubMed Central PMCID: PMC3983093.

6. Kaufman HW, Chen Z, Radcliff J, Batterman HJ, Leake J. Q fever: an under-reported reportable communicable disease. Epidemiol Infect. 2018;146(10):1240–4. Epub 2018/06/26. doi: 10.1017/S0950268818001395. PubMed PMID: 29941056.

7. Georgiev M, Afonso A, Neubauer H, Needham H, Thiery R, Rodolakis A, et al. Q fever in humans and farm animals in four European countries, 1982 to 2010. Euro Surveill. 2013;18(8). Epub 2013/02/21. PubMed PMID: 23449232.

8. Bjork A, Marsden-Haug N, Nett RJ, Kersh GJ, Nicholson W, Gibson D, et al. First reported multistate human Q fever outbreak in the United States, 2011. Vector Borne Zoonotic Dis. 2014;14(2):111-7. Epub 2013/12/18. doi: 10.1089/vbz.2012.1202. PubMed PMID: 24350648.

9. Dijkstra F, van der Hoek W, Wijers N, Schimmer B, Rietveld A, Wijkmans CJ, et al. The 2007–2010 Q fever epidemic in The Netherlands: characteristics of notified acute Q fever patients and the association with dairy goat farming. FEMS Immunol Med Microbiol. 2012;64(1):3-12. doi: 10.1111/j.1574-695X.2011.00876.x. PubMed PMID: 22066649.

10. Madariaga MG, Rezai K, Trenholme GM, Weinstein RA. Q fever: a biological weapon in your backyard. Lancet Infect Dis. 2003;3(11):709–21. doi: 10.1016/s1473-3099(03)00804-1. PubMed PMID: 14592601.

11. Khavkin T, Tabibzadeh SS. Histologic, immunofluorescence, and electron microscopic study of infectious process in mouse lung after intranasal challenge with Coxiella burnetii. Infect Immun. 1988;56(7):1792–9. PubMed PMID: 3290107; PubMed Central PMCID: PMC259479.

12. Stein A, Louveau C, Lepidi H, Ricci F, Baylac P, Davoust B, et al. Q fever pneumonia: virulence of Coxiella burnetii pathovars in a murine model of aerosol infection. Infect Immun. 2005;73(4):2469–77. doi: 10.1128/IAI.73.4.2469-2477.2005. PubMed PMID: 15784593; PubMed Central PMCID: PMC1087393.

13. Heinzen RA, Scidmore MA, Rockey DD, Hackstadt T. Differential interaction with endocytic and exocytic pathways distinguish parasitophorous vacuoles of Coxiella burnetii and Chlamydia trachomatis. Infect Immun. 1996;64(3):796–809. PubMed PMID: 8641784; PubMed Central PMCID: PMC173840.

14. Campoy EM, Zoppino FC, Colombo MI. The early secretory pathway contributes to the growth of the Coxiella-replicative niche. Infect Immun. 2011;79(1):402–13. Epub 2010/10/11. doi: 10.1128/IAI.00688-10. PubMed PMID: 20937765; PubMed Central PMCID: PMC3019900.

15. Romano PS, Gutierrez MG, Berón W, Rabinovitch M, Colombo MI. The autophagic pathway is actively modulated by phase II Coxiella burnetii to efficiently replicate in the host cell. Cell Microbiol. 2007;9(4):891–909. Epub 2006/11/03. doi: 10.1111/j.1462-5822.2006.00838.x. PubMed PMID: 17087732.

16. Voth DE, Heinzen RA. Lounging in a lysosome: the intracellular lifestyle of Coxiella burnetii. Cell Microbiol. 2007;9(4):829–40. doi: 10.1111/j.1462-5822.2007.00901.x. PubMed PMID: 17381428.

17. Coleman SA, Fischer ER, Howe D, Mead DJ, Heinzen RA. Temporal analysis of Coxiella burnetii morphological differentiation. J Bacteriol. 2004;186(21):7344–52. doi: 10.1128/JB.186.21.7344-7352.2004. PubMed PMID: 15489446; PubMed Central PMCID: PMC523218.

18. Moffatt JH, Newton P, Newton HJ. Coxiella burnetii: turning hostility into a home. Cell Microbiol. 2015;17(5):621–31. Epub 2015/03/30. doi: 10.1111/cmi.12432. PubMed PMID: 25728389.

19. Qiu J, Luo ZQ. Legionella and Coxiella effectors: strength in diversity and activity. Nat Rev Microbiol. 2017;15(10):591–605. Epub 2017/07/17. doi: 10.1038/nrmicro.2017.67. PubMed PMID: 28713154.

20. Carey KL, Newton HJ, Lührmann A, Roy CR. The Coxiella burnetii Dot/Icm system delivers a unique repertoire of type IV effectors into host cells and is required for intracellular replication. PLoS Pathog. 2011;7(5):e1002056. Epub 2011/05/26. doi: 10.1371/journal.ppat.1002056. PubMed PMID: 21637816; PubMed Central PMCID: PMC3102713.

21. Newton HJ, Kohler LJ, McDonough JA, Temoche-Diaz M, Crabill E, Hartland EL, et al. A screen of Coxiella burnetii mutants reveals important roles for Dot/Icm effectors and host autophagy in vacuole biogenesis. PLoS Pathog. 2014;10(7):e1004286. Epub 2014/07/31. doi: 10.1371/journal.ppat.1004286. PubMed PMID: 25080348; PubMed Central PMCID: PMC4117601.

22. Beare PA, Gilk SD, Larson CL, Hill J, Stead CM, Omsland A, et al. Dot/Icm type IVB secretion system requirements for Coxiella burnetii growth in human macrophages. mBio. 2011;2(4):e00175–11. Epub 2011/09/01. doi: 10.1128/mBio.00175-11. PubMed PMID: 21862628; PubMed Central PMCID: PMC3163939.

23. Tam JC, Jacques DA. Intracellular immunity: finding the enemy within--how cells recognize and respond to intracellular pathogens. J Leukoc Biol. 2014;96(2):233–44. Epub 2014/06/04. doi: 10.1189/jlb.4RI0214-090R. PubMed PMID: 24899588; PubMed Central PMCID: PMC4192899.

24. Akira S, Uematsu S, Takeuchi O. Pathogen recognition and innate immunity. Cell. 2006;124(4):783-801. doi: 10.1016/j.cell.2006.02.015. PubMed PMID: 16497588.

25. Zamboni DS, Campos MA, Torrecilhas AC, Kiss K, Samuel JE, Golenbock DT, et al. Stimulation of toll-like receptor 2 by Coxiella burnetii is required for macrophage production of pro-inflammatory cytokines and resistance to infection. J Biol Chem. 2004;279(52):54405–15. Epub 2004/10/13. doi: 10.1074/jbc.M410340200. PubMed PMID: 15485838.

26. Meghari S, Honstettre A, Lepidi H, Ryffel B, Raoult D, Mege JL. TLR2 is necessary to inflammatory response in Coxiella burnetii infection. Ann N Y Acad Sci. 2005;1063:161–6. doi: 10.1196/annals.1355.025. PubMed PMID: 16481508.

27. Bradley WP, Boyer MA, Nguyen HT, Birdwell LD, Yu J, Ribeiro JM, et al. Primary Role for Toll-Like Receptor-Driven Tumor Necrosis Factor Rather than Cytosolic Immune Detection in Restricting Coxiella burnetii Phase II Replication within Mouse Macrophages. Infect Immun. 2016;84(4):998–1015. Epub 2016/03/24. doi: 10.1128/IAI.01536-15. PubMed PMID: 26787725; PubMed Central PMCID: PMC4807492.

28. Apostolaki M, Armaka M, Victoratos P, Kollias G. Cellular mechanisms of TNF function in models of inflammation and autoimmunity. Curr Dir Autoimmun. 2010;11:1–26. Epub 2010/02/18. doi: 10.1159/000289195. PubMed PMID: 20173385.

29. Wajant H, Pfizenmaier K, Scheurich P. Tumor necrosis factor signaling. Cell Death Differ. 2003;10(1):45–65. doi: 10.1038/sj.cdd.4401189. PubMed PMID: 12655295.

30. Vassalli P. The pathophysiology of tumor necrosis factors. Annu Rev Immunol. 1992;10:411–52. doi: 10.1146/annurev.iy.10.040192.002211. PubMed PMID: 1590993.

31. Wajant H, Siegmund D. TNFR1 and TNFR2 in the Control of the Life and Death Balance of Macrophages. Front Cell Dev Biol. 2019;7:91. Epub 2019/05/29. doi: 10.3389/fcell.2019.00091. PubMed PMID: 31192209; PubMed Central PMCID: PMC6548990.

32. Ali T, Kaitha S, Mahmood S, Ftesi A, Stone J, Bronze MS. Clinical use of anti-TNF therapy and increased risk of infections. Drug Healthc Patient Saf. 2013;5:79–99. Epub 2013/03/28. doi: 10.2147/DHPS.S28801. PubMed PMID: 23569399; PubMed Central PMCID: PMC3615849.

33. Hirsch J, Astrahan A, Odeh M, Elias N, Rosner I, Rimar D, et al. Q Fever Risk in Patients Treated with Chronic Antitumor Necrosis Factor-Alpha Therapy. Case Rep Infect Dis. 2016;2016:4586150. Epub 2016/08/30. doi: 10.1155/2016/4586150. PubMed PMID: 27656302; PubMed Central PMCID: PMC5021454.

34. Koo S, Marty FM, Baden LR. Infectious complications associated with immunomodulating biologic agents. Hematol Oncol Clin North Am. 2011;25(1):117–38. doi: 10.1016/j.hoc.2010.11.009. PubMed PMID: 21236394.

35. Naujoks J, Tabeling C, Dill BD, Hoffmann C, Brown AS, Kunze M, et al. IFNs Modify the Proteome of Legionella-Containing Vacuoles and Restrict Infection Via IRG1-Derived Itaconic Acid. PLoS Pathog. 2016;12(2):e1005408. Epub 2016/02/01. doi: 10.1371/journal.ppat.1005408. PubMed PMID: 26829557; PubMed Central PMCID: PMC4734697.

36. Chen M, Sun H, Boot M, Shao L, Chang SJ, Wang W, et al. Itaconate is an effector of a Rab GTPase cell-autonomous host defense pathway against. Science. 2020;369(6502):450-5. doi: 10.1126/science.aaz1333. PubMed PMID: 32703879.

37. Hoffmann E, Machelart A, Belhaouane I, Deboosere N, Pauwels A-M, Saint-Andre J-P, et al. IRG1 controls immunometabolic host response and restricts intracellular Mycobacterium tuberculosis infection2019.

38. Beare PA, Howe D, Cockrell DC, Omsland A, Hansen B, Heinzen RA. Characterization of a Coxiella burnetii ftsZ mutant generated by Himar1 transposon mutagenesis. J Bacteriol. 2009;191(5):1369–81. Epub 2008/12/29. doi: 10.1128/JB.01580-08. PubMed PMID: 19114492; PubMed Central PMCID: PMC2648191.

39. Omsland A, Beare PA, Hill J, Cockrell DC, Howe D, Hansen B, et al. Isolation from animal tissue and genetic transformation of Coxiella burnetii are facilitated by an improved axenic growth medium. Appl Environ Microbiol. 2011;77(11):3720–5. Epub 2011/04/08. doi: 10.1128/AEM.02826-10. PubMed PMID: 21478315; PubMed Central PMCID: PMC3127619.

40. Livak KJ, Schmittgen TD. Analysis of relative gene expression data using real-time quantitative PCR and the 2(-Delta Delta C(T)) Method. Methods. 2001;25(4):402–8. doi: 10.1006/meth.2001.1262. PubMed PMID: 11846609.

41. Chen G, Goeddel DV. TNF-R1 signaling: a beautiful pathway. Science. 2002;296(5573):1634-5. doi: 10.1126/science.1071924. PubMed PMID: 12040173.

42. Blander JM. A long-awaited merger of the pathways mediating host defence and programmed cell death. Nat Rev Immunol. 2014;14(9):601–18. doi: 10.1038/nri3720. PubMed PMID: 25145756.

43. Peltzer N, Darding M, Walczak H. Holding RIPK1 on the Ubiquitin Leash in TNFR1 Signaling. Trends Cell Biol. 2016;26(6):445–61. Epub 2016/02/11. doi: 10.1016/j.tcb.2016.01.006. PubMed PMID: 26877205.

44. Feoktistova M, Leverkus M. Programmed necrosis and necroptosis signalling. FEBS J. 2015;282(1):19-31. Epub 2014/11/11. doi: 10.1111/febs.13120. PubMed PMID: 25327580.

45. Tummers B, Green DR. Caspase-8: regulating life and death. Immunol Rev. 2017;277(1):76–89. doi: 10.1111/imr.12541. PubMed PMID: 28462525; PubMed Central PMCID: PMC5417704.

46. Kaiser WJ, Upton JW, Long AB, Livingston-Rosanoff D, Daley-Bauer LP, Hakem R, et al. RIP3 mediates the embryonic lethality of caspase-8-deficient mice. Nature. 2011;471(7338):368-72. Epub 2011/03/02. doi: 10.1038/nature09857. PubMed PMID: 21368762; PubMed Central PMCID: PMC3060292.

47. Berger SB, Kasparcova V, Hoffman S, Swift B, Dare L, Schaeffer M, et al. Cutting Edge: RIP1 kinase activity is dispensable for normal development but is a key regulator of inflammation in SHARPIN-deficient mice. J Immunol. 2014;192(12):5476–80. Epub 20140512. doi: 10.4049/jimmunol.1400499. PubMed PMID: 24821972; PubMed Central PMCID: PMC4048763.

48. Xie QW, Kashiwabara Y, Nathan C. Role of transcription factor NF-kappa B/Rel in induction of nitric oxide synthase. J Biol Chem. 1994;269(7):4705–8. PubMed PMID: 7508926.

49. Chakravortty D, Hensel M. Inducible nitric oxide synthase and control of intracellular bacterial pathogens. Microbes Infect. 2003;5(7):621–7. doi: 10.1016/s1286-4579(03)00096-0. PubMed PMID: 12787738.

50. Brennan RE, Russell K, Zhang G, Samuel JE. Both inducible nitric oxide synthase and NADPH oxidase contribute to the control of virulent phase I Coxiella burnetii infections. Infect Immun. 2004;72(11):6666–75. doi: 10.1128/IAI.72.11.6666-6675.2004. PubMed PMID: 15501800; PubMed Central PMCID: PMC523001.

51. Howe D, Barrows LF, Lindstrom NM, Heinzen RA. Nitric oxide inhibits Coxiella burnetii replication and parasitophorous vacuole maturation. Infect Immun. 2002;70(9):5140–7. doi: 10.1128/iai.70.9.5140-5147.2002. PubMed PMID: 12183564; PubMed Central PMCID: PMC128226.

52. Lambeth JD. NOX enzymes and the biology of reactive oxygen. Nat Rev Immunol. 2004;4(3):181–9. doi: 10.1038/nri1312. PubMed PMID: 15039755.

53. Blaser H, Dostert C, Mak TW, Brenner D. TNF and ROS Crosstalk in Inflammation. Trends Cell Biol. 2016;26(4):249–61. Epub 2016/01/12. doi: 10.1016/j.tcb.2015.12.002. PubMed PMID: 26791157.

54. Ivashkiv LB, Donlin LT. Regulation of type I interferon responses. Nat Rev Immunol. 2014;14(1):36–49. doi: 10.1038/nri3581. PubMed PMID: 24362405; PubMed Central PMCID: PMC4084561.

55. Honda K, Taniguchi T. IRFs: master regulators of signalling by Toll-like receptors and cytosolic pattern-recognition receptors. Nat Rev Immunol. 2006;6(9):644–58. doi: 10.1038/nri1900. PubMed PMID: 16932750.

56. Wang Z, Choi MK, Ban T, Yanai H, Negishi H, Lu Y, et al. Regulation of innate immune responses by DAI (DLM-1/ZBP1) and other DNA-sensing molecules. Proc Natl Acad Sci U S A. 2008;105(14):5477–82. Epub 2008/03/28. doi: 10.1073/pnas.0801295105. PubMed PMID: 18375758; PubMed Central PMCID: PMC2291118.

57. Yarilina A, Ivashkiv LB. Type I interferon: a new player in TNF signaling. Curr Dir Autoimmun. 2010;11:94–104. Epub 2010/02/18. doi: 10.1159/000289199. PubMed PMID: 20173389; PubMed Central PMCID: PMC2827816.

58. Degrandi D, Hoffmann R, Beuter-Gunia C, Pfeffer K. The proinflammatory cytokine-induced IRG1 protein associates with mitochondria. J Interferon Cytokine Res. 2009;29(1):55–67. doi: 10.1089/jir.2008.0013. PubMed PMID: 19014335.

59. Michelucci A, Cordes T, Ghelfi J, Pailot A, Reiling N, Goldmann O, et al. Immune-responsive gene 1 protein links metabolism to immunity by catalyzing itaconic acid production. Proc Natl Acad Sci U S A. 2013;110(19):7820–5. Epub 2013/04/22. doi: 10.1073/pnas.1218599110. PubMed PMID: 23610393; PubMed Central PMCID: PMC3651434.

60. Price JV, Russo D, Ji DX, Chavez RA, DiPeso L, Lee AY, et al. IRG1 and Inducible Nitric Oxide Synthase Act Redundantly with Other Interferon-Gamma-Induced Factors To Restrict Intracellular Replication of Legionella pneumophila. mBio. 2019;10(6). Epub 2019/11/12. doi: 10.1128/mBio.02629-19. PubMed PMID: 31719183; PubMed Central PMCID: PMC6851286.

61. McKinney JD, Höner zu Bentrup K, Muñoz-Elías EJ, Miczak A, Chen B, Chan WT, et al. Persistence of Mycobacterium tuberculosis in macrophages and mice requires the glyoxylate shunt enzyme isocitrate lyase. Nature. 2000;406(6797):735-8. doi: 10.1038/35021074. PubMed PMID: 10963599.

62. van Schaik EJ, Tom M, Woods DE. Burkholderia pseudomallei isocitrate lyase is a persistence factor in pulmonary melioidosis: implications for the development of isocitrate lyase inhibitors as novel antimicrobials. Infect Immun. 2009;77(10):4275–83. Epub 2009/07/20. doi: 10.1128/IAI.00609-09. PubMed PMID: 19620343; PubMed Central PMCID: PMC2747945.

63. Hedges JF, Robison A, Kimmel E, Christensen K, Lucas E, Ramstead A, et al. Type I Interferon Counters or Promotes Coxiella burnetii Replication Dependent on Tissue. Infect Immun. 2016;84(6):1815–25. Epub 2016/05/24. doi: 10.1128/IAI.01540-15. PubMed PMID: 27068091; PubMed Central PMCID: PMC4907146.

64. Mestan J, Brockhaus M, Kirchner H, Jacobsen H. Antiviral activity of tumour necrosis factor. Synergism with interferons and induction of oligo-2’,5’-adenylate synthetase. J Gen Virol. 1988;69 (Pt 12):3113-20. doi: 10.1099/0022-1317-69-12-3113. PubMed PMID: 2462015.

65. Fink K, Martin L, Mukawera E, Chartier S, De Deken X, Brochiero E, et al. IFNβ/TNFα synergism induces a non-canonical STAT2/IRF9-dependent pathway triggering a novel DUOX2 NADPH oxidase-mediated airway antiviral response. Cell Res. 2013;23(5):673–90. Epub 2013/04/02. doi: 10.1038/cr.2013.47. PubMed PMID: 23545780; PubMed Central PMCID: PMC3641604.

66. Mancuso G, Midiri A, Biondo C, Beninati C, Zummo S, Galbo R, et al. Type I IFN signaling is crucial for host resistance against different species of pathogenic bacteria. J Immunol. 2007;178(5):3126–33. doi: 10.4049/jimmunol.178.5.3126. PubMed PMID: 17312160.

67. Daniels BP, Kofman SB, Smith JR, Norris GT, Snyder AG, Kolb JP, et al. The Nucleotide Sensor ZBP1 and Kinase RIPK3 Induce the Enzyme IRG1 to Promote an Antiviral Metabolic State in Neurons. Immunity. 2019;50(1):64–76.e4. Epub 2019/01/08. doi: 10.1016/j.immuni.2018.11.017. PubMed PMID: 30635240; PubMed Central PMCID: PMC6342485.

68. Kohl L, Siddique MNAA, Bodendorfer B, Berger R, Preikschat A, Daniel C, et al. Macrophages inhibit Coxiella burnetii by the ACOD1-itaconate pathway for containment of Q fever. EMBO Mol Med. 2023;15(2):e15931. Epub 20221207. doi: 10.15252/emmm.202215931. PubMed PMID: 36479617; PubMed Central PMCID: PMC9906395.

69. McFadden BA, Purohit S. Itaconate, an isocitrate lyase-directed inhibitor in Pseudomonas indigofera. J Bacteriol. 1977;131(1):136–44. PubMed PMID: 17593; PubMed Central PMCID: PMC235402.

70. Patel TR, McFadden BA. Caenorhabditis elegans and Ascaris suum: inhibition of isocitrate lyase by itaconate. Exp Parasitol. 1978;44(2):262–8. doi: 10.1016/0014-4894(78)90107-8. PubMed PMID: 658222.

71. Berg IA, Filatova LV, Ivanovsky RN. Inhibition of acetate and propionate assimilation by itaconate via propionyl-CoA carboxylase in isocitrate lyase-negative purple bacterium Rhodospirillum rubrum. FEMS Microbiol Lett. 2002;216(1):49–54. doi: 10.1111/j.1574-6968.2002.tb11413.x. PubMed PMID: 12423751.

72. Cordsmeier A, Wagner N, Lührmann A, Berens C. Defying Death - How. Yale J Biol Med. 2019;92(4):619–28. Epub 2019/12/20. PubMed PMID: 31866777; PubMed Central PMCID: PMC6913804.

73. Lührmann A, Nogueira CV, Carey KL, Roy CR. Inhibition of pathogen-induced apoptosis by a Coxiella burnetii type IV effector protein. Proc Natl Acad Sci U S A. 2010;107(44):18997–9001. Epub 2010/10/13. doi: 10.1073/pnas.1004380107. PubMed PMID: 20944063; PubMed Central PMCID: PMC2973885.

74. Klingenbeck L, Eckart RA, Berens C, Lührmann A. The Coxiella burnetii type IV secretion system substrate CaeB inhibits intrinsic apoptosis at the mitochondrial level. Cell Microbiol. 2013;15(4):675–87. Epub 2012/11/27. doi: 10.1111/cmi.12066. PubMed PMID: 23126667.

75. Bisle S, Klingenbeck L, Borges V, Sobotta K, Schulze-Luehrmann J, Menge C, et al. The inhibition of the apoptosis pathway by the Coxiella burnetii effector protein CaeA requires the EK repetition motif, but is independent of survivin. Virulence. 2016;7(4):400–12. Epub 2016/01/13. doi: 10.1080/21505594.2016.1139280. PubMed PMID: 26760129; PubMed Central PMCID: PMC4871633.

76. Pollock TY, Vázquez Marrero VR, Brodsky IE, Shin S. TNF licenses macrophages to undergo rapid caspase-1, -11, and -8-mediated cell death that restricts Legionella pneumophila infection. PLoS Pathog. 2023;19(6):e1010767. Epub 20230606. doi: 10.1371/journal.ppat.1010767. PubMed PMID: 37279255; PubMed Central PMCID: PMC10275475.

